# scAbsolute: measuring single-cell ploidy and replication status

**DOI:** 10.1101/2022.11.14.516440

**Authors:** Michael P. Schneider, Amy Cullen, Justina Pangonyte, Jason Skelton, Harvey Major, Elke Van Oudenhove, Maria J. Garcia, Blas Chaves-Urbano, Anna M. Piskorz, James D. Brenton, Geoff Macintyre, Florian Markowetz

## Abstract

Cancer cells often exhibit DNA copy number aberrations and can vary widely in their ploidy. Correct estimation of the ploidy of single cell genomes is paramount for downstream analysis. Based only on single-cell DNA sequencing information, *scAbsolute* achieves accurate and unbiased measurement of single-cell ploidy and replication status, including whole-genome duplications. We demonstrate *scAbsolute’s* capabilities using experimental cell multiplets, a FUCCI cell cycle expression system, and a benchmark against state-of-the-art methods. *scAbsolute* provides a robust foundation for single-cell DNA sequencing analysis across different technologies and has the potential to enable improvements in a number of downstream analyses.

## Background

Many common cancers are characterised by chromosomal instability (CIN) and as a result, extensive copy number aberrations (CNAs) [1, 2]. CNAs alter the number of copies of genomic regions in a cell, thus creating a background of genomic variation on which selection can act [3]. CNAs can act as drivers of cancer evolution [4–6] and be used to infer phylogenies [7, 8]. Importantly, CNAs have been shown to play a crucial role in cancer treatment and prognosis [9–11], and they correlate with markers of immune evasion and increased activity in proliferation pathways [6, 12, 13]. As a consequence of CIN, tumour cells, unlike most normal cells, substantially vary in the amount of DNA they contain, the so-called ploidy.

Large amounts of DNA in a cell or high levels of ploidy are often associated with whole genome doubling (WGD) [14]. Previous research has shown how WGD can fuel CIN through abnormal mitosis [15–18]. Cells that have undergone WGD can be genomically unstable and tend to accumulate CNAs more quickly, partly because they appear to be able to better cope with the negative effects of deleterious mutations and ongoing CIN. [14, 19–22]. WGD is a common event across cancers [1, 23, 24] and is associated with poor prognosis [25]. At the same time, fundamental questions are still unanswered, for example, the interplay between WGD and CIN in the early stages of tumourigenesis [16, 26].

WGD and CIN are central topics in cancer genomics, and analyses based on single-cell DNA sequencing hold promises for better understanding how these processes shape the landscape of somatic mutations in cancer and how they impact tumour development at the earliest stages [27]. Single-cell genomics offers a much more fine-grained clonal decomposition of malignant tissues and the ability to study properties of rare or small cell population genotypes [28]. Advances in single-cell sequencing technologies [28–34] make it possible to measure CNAs in individual cancer cells. Early technologies relied on whole-genome amplification, an example included in this study is the 10X Genomics Single Cell CNV[35] platform using a commercial microdroplet platform. Several publicly available data sets were produced by this technology which is no longer commercially available. Recently, this approach has been challenged by protocols developed for shallow single-cell WGS, including Direct Library Preparation [28, 31] (DLP/DLP+) and Acoustic Cell Tagmentation [34] (ACT). These two technologies rely on whole-genome single-cell library preparation without amplification (preamplification-free) and single-molecule indexing via direct tagmentation. By using a direct tagmentation step to incorporate index barcodes and sequencing adaptors before subsequent PCR cycles, it is possible to link all original reads to the original single-cell molecules and computationally filter PCR duplicates leading to a more even coverage across the genome [31].

Further progress, however, is held back by the lack of accurate methods to identify the ploidy of single cells, which is a crucial prerequisite for many downstream applications, such as quantifying intratumour heterogeneity and phylogenetic reconstruction of tumour evolution, and also heavily impacts single-nucleotide variation (SNV) detection [36–38]. The challenges of ploidy calling in single-cell data are further aggravated by the fact that different cell cycle states lead to different overall DNA content and introduce spurious copy number changes [39, 40]. As a result, separating cells undergoing DNA replication in S phase from cells in G1/G2 phase is important to reliably measure copy number status across cells.

To address these challenges, we introduce *scAbsolute*, a computational method specifically targeted at inferring individual cells’ ploidy and replication status based on shallow single-cell DNA sequencing data alone. We demonstrate the feasibility of distinguishing cells in different, previously unidentifiable ploidy states, including cells directly after undergoing WGD.

Our research improves on existing models for ploidy estimation across different data types. For bulk sequencing data, many tools aim to estimate tumour purity and ploidy, and identify subclonal copy-number status [23, 37, 41–48], but the problem is challenging, because it is underdetermined with multiple mathematically equivalent solutions existing [23]. Existing approaches for single-cell sequencing data either build on computational steps originally developed for bulk data or use additional experimental information. Most approaches are unable to distinguish between different ploidy solutions in the absence of odd or intermediate copy number states in the data [49]. In practice, tools such as HMMCopy [50] are commonly used [31, 32, 51, 52] and serve as the basis for some novel copy number callers [53, 54]. Ginkgo [55] shows improved performance for calling of accurate ploidy on a single-cell level [56] in simulated data. SCOPE [57] uses normal cells in a cross-sample normalisation step to estimate bin-specific noise terms. However, normal cells are not available for many single-cell sequencing studies and cell lines, limiting its applicability in practice. In a further advance, CHISEL [58] estimates haplotype-specific copy numbers based on B-allele frequency (BAF) estimated across 100s and 1000s of cells. CHISEL requires to pool a large number of reads together, either by increased sequencing depth or by number of cells, and needs a matched normal sample or a sufficient number of normal single cells sequenced. As a BAF-based approach, CHISEL cannot detect recent WGD, because of the need for sufficient number of mutations to arise across different alleles. Alternatively, ploidy information can be inferred experimentally. A common approach is using DAPI fluorescence staining and subsequent Fluorescence-activated Cell Sorting (FACS) [34]. However, this approach requires a ploidy control and a sufficient number of cells as input material, and might introduce a bias in the a-priori selection of cells based on their ploidy profile. Laks *et. al*. [28] suggest that there is a relationship between ploidy and cell size, as observed via microscopy. However, it is unclear to what extent this can be reliably used to determine absolute cell ploidy.

Previous attempts for identifying replication status have been based on features derived in homogeneous samples or limited training sets [28, 59, 60], and it is unclear how well these methods generalise to novel, previously unseen samples or heterogeneous populations. We introduce a purely computational approach that generalises to novel cell populations without requiring new training data, and is more accurate than existing experimental evidence based on FACS sorting of DAPI stained cell populations.

## Results

### The scAbsolute algorithm for calling absolute copy number in single-cell DNA sequencing data

*scAbsolute* aims to find a transform to convert values from a scale of read counts per bin to a scale of absolute copy number.

For a cell with its genome split into *M* fixed-size genomic bins (by default 500 kb), we refer to the (unknown) copy number as *c*_*j*_ and the observed per-bin read count as *x*_*j*_ for each bin *j*. We aim to estimate a scaling factor *ρ*, so that we can directly measure the ploidy *p* of a cell:

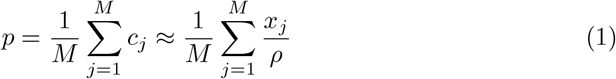

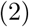

Estimating the value of *ρ* is equivalent to determining the correct copy number states *c*_*j*_, assuming we have a segmented copy number profile. The factor *ρ* denotes the average reads per copy per bin and is a measure of per-cell read coverage that we find useful to compare cells even with very different copy number profiles. The value of *ρ* is a direct measure of the difference in expected mean read counts between neighbouring copy number states. Alternatively, one can imagine the raw read counts clustering around integer multiples of *ρ*, as shown in Fig. 1. Note, that our definition of ploidy is directly proportional to the amount of DNA in a cell, and is not referring to the number of chromosomes in a cell.

**Fig. 1:**
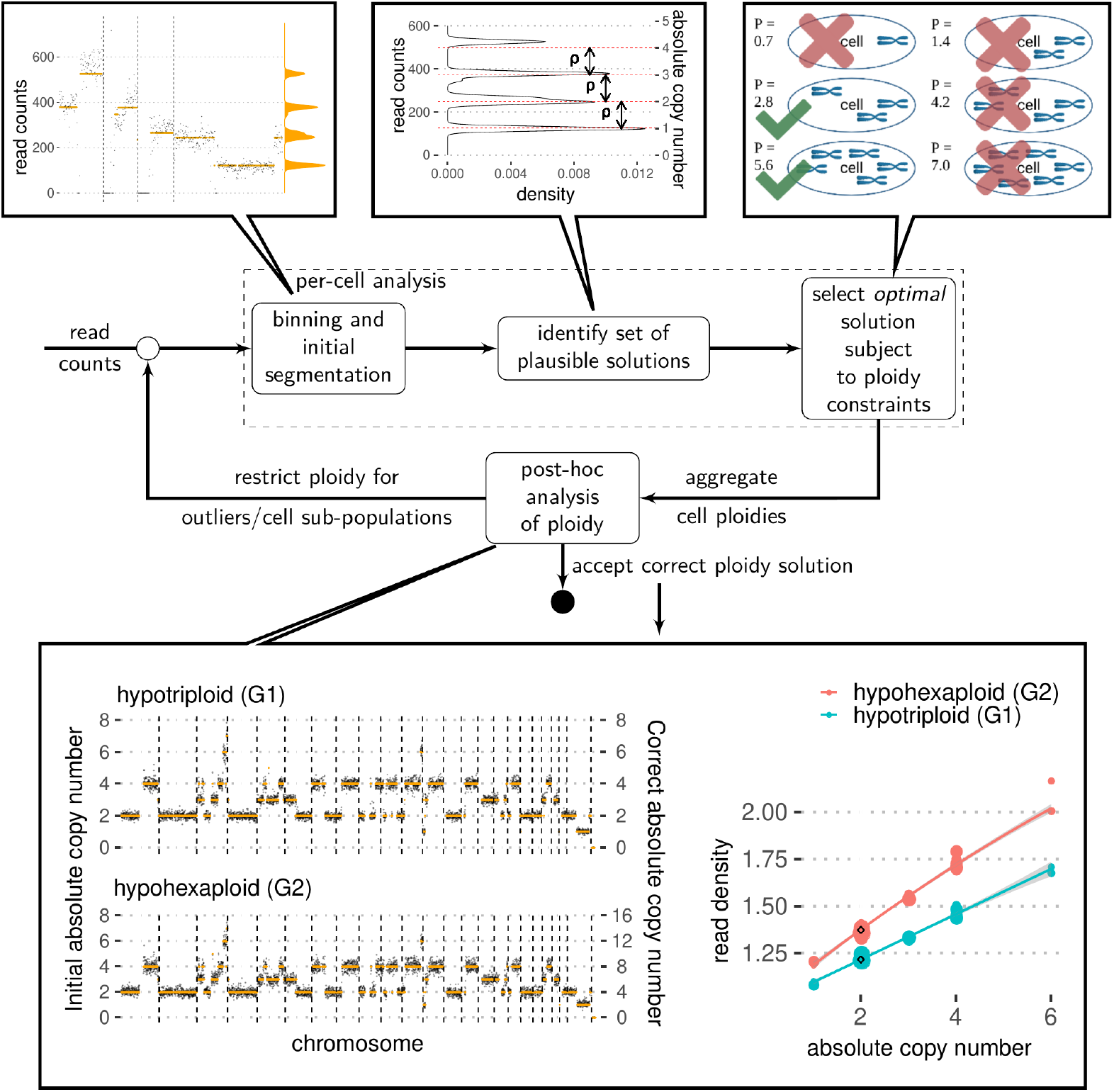
Schematic overview of *scAbsolute* approach. Initially, raw read counts are binned and segmented. The marginal distribution of the segmented read counts is fitted with a constrained Dirichlet Process Gaussian Mixture Model (DPGMM) using stochastic variational inference in order to identify a limited set of plausible solutions. We do this by identifying a constant *ρ*, that converts the scale from a read count to an absolute copy number scale. There exist a set of values of *ρ* that lead to equally possible ploidy solutions (in this case hypotriploid or hypohexaploid solutions). We select the solution with minimum ploidy value subject to per-cell ploidy constraints. Post-hoc analysis of ploidy allows to distinguish previously indistinguishable cell populations based on read density, overcoming the limitation of the above approach. The example shows two (exemplary) copy number profiles for cells in different ploidy states, that have previously been indistinguishable based on copy number profiles alone. We demonstrate that cell ploidy can be determined in the absence of differentiating copy number states and other experimental information, based on per-cell read density alone.

*scAbsolute* directly works on aligned bam files (we support Human and Mouse genome builds) and produces absolute copy number calls across genomic bins. Ploidy and cell-cycle inference takes about 3-10 min per cell (at 500 kb bin resolution), and can be easily parallelized across cells. *scAbsolute* proceeds in a series of steps summarised below (and in Fig. 1); further details can be found in the Methods section.

Step 1: We use a dynamic programming approach based on the PELT algorithm for change point detection [61] with a negative binomial likelihood to find an initial segmentation of our read counts. It is possible to use other segmentation algorithms. Our focus here is not on obtaining a highly accurate segmentation, but rather an initial segmentation that allows us to identify an accurate ploidy estimate.

Step 2: We then consider the marginal distribution of the segmented read counts, i.e. we eliminate the influence of genomic location via segmentation and look at the distribution of the segmented values. Because we are working with a single cell, we know that all copy number states occur only at integer levels. We use a constrained Dirichlet Process Gaussian Mixture Model to fit a mixture of Gaussians to the marginal distributions. The constraint forces the distance between the means of the Gaussians to be constant, analogous to a 1-D grid. The width of the grid then corresponds to a multiple of the scaling factor *ρ*, since we identify the peaks with the discrete copy number states that are scaled by the per-cell read depth.

Step 3: We further constrain the per-cell ploidy solution by choosing a multiple of the scaling factor as the correct solution, based on the resulting cell ploidy. Here, we use the solution with minimum root mean-squared error among all these solutions within the given ploidy window (by default 1.1-8.0). Following this procedure, some of these cells will be fit at the correct ploidy, whereas others may not be (due to the non-identifiability introduced via WGD or G2 phase, see Fig. S1)

Step 4: In order to identify cells in which step 3 leads to an incorrect ploidy assignment - such as G2 phase or WGD cells, or other outliers - we compute the genomic read density given an inferred ploidy in a post-processing step. Deviations from the empirically expected read density at the estimated ploidy indicate an incorrect ploidy solution. If any outliers have been detected in this step, they can be refitted by re-applying steps 1-3, with an updated cell-specific ploidy window.

Below, we present the *scAbsolute* method with three applications: identifying previously unidentifiable ploidy cases, identifying cells undergoing DNA replication, and a comparison of *scAbsolute* with existing methods for determining per-cell ploidy.

### Accurate ploidy estimation based on read density

*scAbsolute* addresses the ploidy identifiability problem using an approach based on read density. This innovation is encompassed in step 4 of the algorithm. The approach is applicable to all single-cell DNA sequencing technologies that do not use whole-genome amplification and have a sufficient per-cell read coverage (at least *ρ* = 75 at 500 kb resolution with paired-end reads; corresponding to a coverage of about 0.01-0.02x or about a million reads for a normal, diploid, human cell). Single-end read data would require a much higher read depth to detect a sufficient number of overlapping reads to make our method applicable. Without a whole-genome amplification step we can directly reference reads to physical copies of the genome, and this provides the necessary constraint to uniquely identify a ploidy solution. We use the start and end position of a given read pair, and measure how many other reads overlap with it. Using the fact that in the absence of a whole-genome amplification step, we expect the number of overlapping reads to be limited by the number of physical copies of the molecules, we can use this to build a model of how many overlapping reads there are on average in any genomic bin (see Fig. S2). To make this approach sufficiently robust and generalisable, we use a genome-wide measure of read density, i.e. the number of overlapping reads per region of the genome, to create a reference distribution of the expected mean number of overlapping reads. Importantly, this measure of read density depends on two variables, only: an individual cell’s ploidy and the read depth per cell as captured by *ρ*.

First, we obtain a ploidy-normalized per-cell value of read density, by regressing the observed read density across copy number states and chromosomes, and predicting an expected value for a copy number state of 2 (see Fig. 1, bottom panel). This allows us to make a prediction independently of varying copy number states in different cells and even in the absence of a copy number state of 2. Lastly, we need to account for varying per-cell read depths (as measured by the *ρ* value). We normalize the per-cell value, by fitting a simple quadratic function to the observed read densities (Figs. 2a and 2b). This makes it possible to directly compare a new cell to the predicted value of read density and observe any strong deviations on the level of the residuals. The model is currently strongest in the range of 75 *−*150 *ρ*, as we do have substantially more cells at this read depth and generally less reliable for higher values of *ρ*. However, more overlapping reads are detected for higher values of *ρ*, making the distinction between different ploidy cases easier for higher values of *ρ* in general (Figs. 2b and 2d). Here, we make the assumption that the majority of cells in the reference set are in G1 phase, and we correctly identify the individual per-cell ploidies for the majority of these cells. Another issue in real world data is the propensity of observing higher read depth for cells with higher ploidies. In practice these high ploidy cells are often cells in G2 phase or S phase, additionally introducing a bias in the model for higher values of *ρ*.

**Fig. 2:**
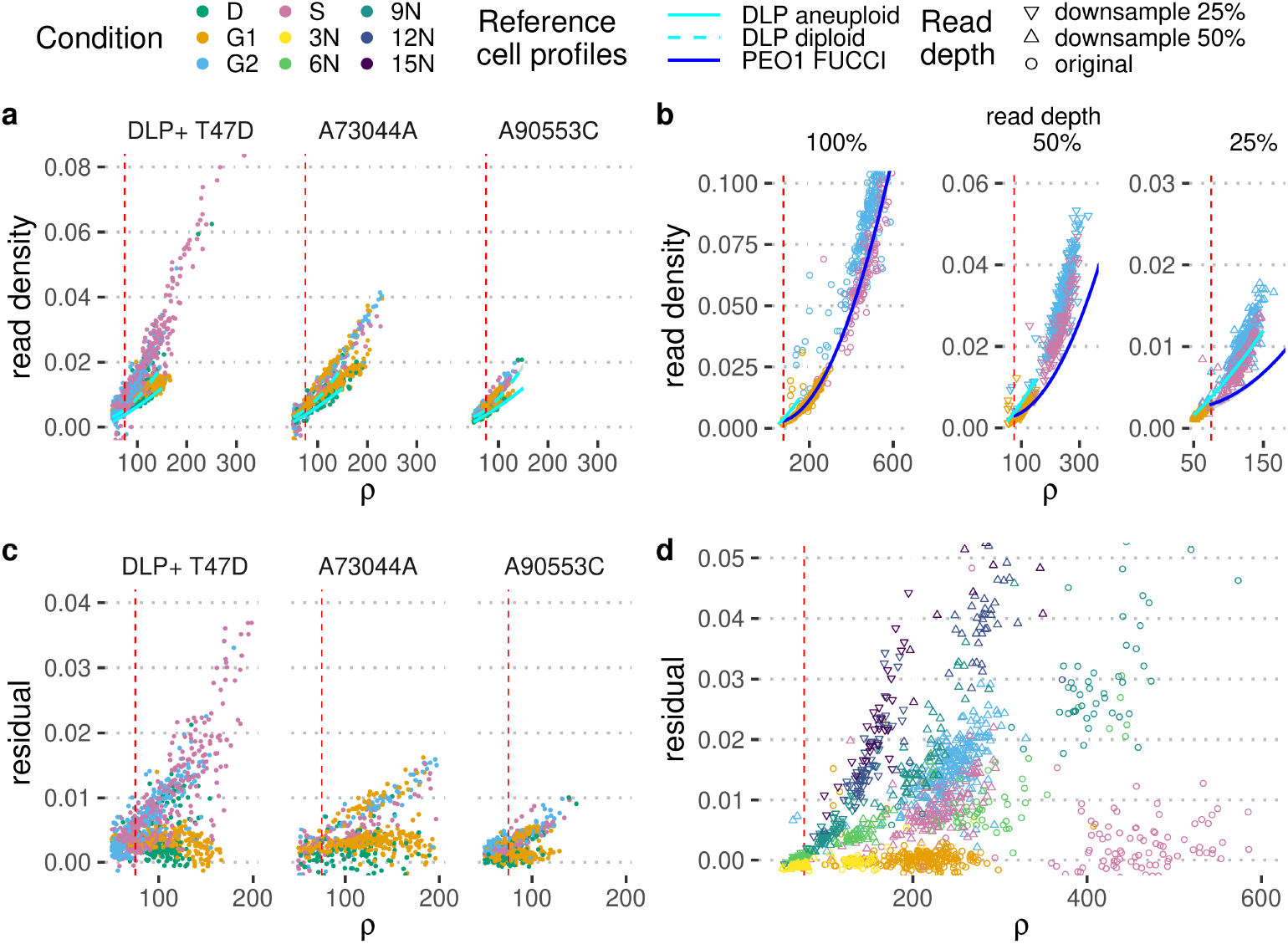
Accurate ploidy estimation based on read density. **a**) Read depth *ρ* versus observed read density for cells sorted into different cell cycle stages (apoptotic (D), G1, G2, and S phase) via FACS using DAPI staining. Each panel represents a different library: T-47D is a cancer cell line, and A73044A and A90553C are libraries consisting of normal cells (GM18507 cell line/SA928). Models of expected read density for a given read depth *ρ* based on a reference populations of cells sequenced with DLP+ are indicated in cyan. *ρ* = 75 is indicated by the red dashed line. **b**) Read depth *ρ* versus observed read density for PEO1 cells sorted into different cell cycle stages (apoptotic (D), G1, G2, and S phase) via the FUCCI cell cycle marker system. Each panel represents a systematic downsampling of cells (100%, and to 50% and 25% of original read depth). A model of expected read density for a given read depth *ρ* based on a reference populations of cells in G1 and S phase sequenced with mDLP+ is indicated in blue, and a model from **a** is included for reference. **c**) Read depth *ρ* versus residual of the observed read density compared to the expected from the trained model in **a** for cells sorted into different cell cycle stages via FACS using DAPI staining. **d**) Read depth *ρ* versus residual of the observed read density compared to the expected from the trained model in **b** (PEO1 FUCCI) for cells sorted into different cell cycle stages via FUCCI cell cycle marker system. Additionally, residuals for cell multiplets are included. Cell multiplets are colour coded by ploidy state as determined experimentally via mixing of cell multiplets containing 1, 2, 3, 4 or 5 individual cells representing 3N, 6N, 9N, 12N, and 15N ploidy, within a single well prior to library preparation and sequencing. Cell multiplets and FUCCI cells are downsampled to 25% and 50% of original read depth to be included where appropriate.

To show that this approach generalizes across data sets, we fit a model to a series of high-depth (with *ρ >* 75) DLP+ data sets, holding out three single-cell libraries: One library based on the T-47D sample with 196 cells in G1, and 154 cells in G2 phase respectively, and two libraries from SA928 (normal cell line) with 664 cells in G1, and 362 cells in G2 phase, respectively. We observe a clear difference in read densities between cells in G1 and G2 phase of the cell cycle, with S phase cells taking a somewhat intermediate position, with some cells more closely aligned to G2 cells possibly corresponding to late-replicating S phase cells, and others to early-replicating cells in S phase. Using the hold-out set, we accurately classify 89% of cells in T-47D, and about 66% of cells in SA928 when looking at cells with a read depth of *ρ* ≥75. Note that in the later case, we can see clear evidence for potential outlier cells, indicating that this might be an underestimate of actual predictive performance, due to the challenges with FACS using DAPI staining based gating of cell populations to distinguish cells in different cell cycle stages (see also Figs. S3 and S4 and the discussion of the T-47D sample in the next section). However, we also note that prediction in the case of normal, diploid cells is somewhat more difficult, as the per-cell regression in these cases is impacted more strongly by outliers.

To address the difficulty of clearly separating cells in different cell cycle stages with FACS sorting using DAPI staining, and to show the generalisability of our approach, we transduced ovarian cancer PEO1-STOP cells with the FUCCI expression system facilitating accurate sorting of cells based on markers indicating different phases of the cell cycle including Geminin, expressed in S and G2 phase, and Cdt1, expressed in G1 and early S phase. We then performed single cell DNA sequencing of 288 cells in each phase of the cell cycle (G1, S and G2). In addition, to generate known, variable ploidy states for the same cell line, we plated multiple cells into individual wells and treated each well as a ‘single cell’ when performing single-cell DNA sequencing. We sequenced 288 of these artificial multiplet cells, and as the PEO1 line is approximately triploid these multiplets represented known ploidy states of 3N, 6N, 9N, 12N, and 15N cells, based on the number of cells per multiplet. The artificial multiplets were experimentally categorised as G1 using negative selection against Geminin (Geminin is not expressed in G1 phase, but expressed from the transition from G1 to S phase onwards) antibody staining. For each condition we obtained 64 cells, except for 10N where we have 32 cells. We initially focused our analysis on those cells that are assigned identical ploidy corresponding to 3N/G1 cells after the first pass of *scAbsolute* (some of the cells were correctly assigned higher ploidies because of individual copy number aberrations). We used the G1 and S phase cells from the FUCCI experiment to create a model of expected read density, and saw a clear deviation for cells in G2 phase (Fig. 2b). Similarly, artificial cell multiplets (6N) cells showed comparatively high levels of read density given their read depth *ρ*. We compared the FUCCI cells downsampled to 50% read depth, and could distinguish G1 from G2/S phase cells with 97% accuracy at a value of *ρ*≈ 200 (Fig. 2d). Looking at the artificial cell multiplets, we could distinguish G1 cells from cells with higher ploidy (6N-12N) with a similar high accuracy of 97%, both in the case of downsampled data (50% downsampled, *ρ* ≈200), and for the original data (*ρ*≈ 400) using differences in the residual between observed and expected read densities. There was a clear trend for higher ploidies (6N, 9N, 12N, 15N) to have higher read densities and associated with this, a higher residual (Fig. 2d). The cells appeared very similar on the level of per-cell copy number profiles and could not be distinguished on their basis alone (see Fig. S5).

### Identifying cells undergoing DNA replication

Cells undergoing DNA replication exhibit temporary CNAs as a consequence of different regions of the genome replicating at different time points [39, 40]. In order to obtain a correct understanding of what constitutes actual tumour heterogeneity at the copy number level, and what are just ephemeral CNAs as a consequence of replication activity, it is therefore necessary to correctly identify cells undergoing replication.

To achieve this goal, we introduce a basic measure of cycling activity based on fixed-width genomic bin counts and their respective replication time. This measure is computed on a per-cell basis after the initial ploidy fit, but the measure does not depend on the ploidy estimate itself. We measure for each copy number segment in a cell whether we can observe a statistically significant trend in the observed counts as a function of replication time, correcting for the joint effect of GC content and mappability using the partial correlation trend test[62]. Replication time information is based on information from the ENCODE project. We define the cycling activity measure as the median of the per-segment trend test statistic weighted by the segment lengths. The advantages of this measure are: i) applicability across sequencing technologies (10X, DLP, ACT) ii) robustness across a range and mix of copy number profiles (as expected in a complex tumour sample) iii) unsupervised method (no training data required).

Using only four features (per-cell read depth, a measure of overdispersion, cycling activity, and a measure of mutual information between reads and replication state of a bin), we trained a random forest classifier to identify S-phase cells and achieved an accuracy of 92%. This analysis was performed using the same data that was used in a previous study, Laks *et. al*. [28] where a complex cell-feature based classifier achieved 90% accuracy. Using only the cycling activity and the mutual information measure in an unsupervised manner, we achieved 86% accuracy. Performance using only this single feature classifier was different between normal cells (DLP-SA928) at 90% accuracy and T-47D human breast cancer cell line (T-47D) at 78% accuracy. However, we believed this to be an underestimate of real performance, due to potentially erroneous assignments to cell cycle stages by FACS based sorting; a closer look at the underlying classification of S phase cells indicated that there were two distinct populations classified as S phase based on FACS (Figs. S3, 3b and 3c). By looking at the copy number profiles, we saw that the cells classified as S phase according to FACS and G1 phase according to our classifier appeared to be closer to the G1 phase cells based on a UMAP representation of the copy number profiles (Figs. S3 and S4). Similarly, by looking at the raw copy number profiles, there appeared to be two different subpopulations among the cells lumped together as S phase cells using FACS. This difficulty of gating cells in different phases of the cell cycle using FACS and DAPI staining, was supported in our own data by the addition of Geminin staining to control for cell cycle, showing that DAPI staining based selection led to the inclusion of a relatively large proportion of cells in S phase (see Fig. S6), if the gating was not conducted very carefully.

To more clearly separate cells in different stages of the cell cycle and to address the limitation of sorting cells with FACS using DAPI staining for cell cycle annotation, we used cells transfected with the FUCCI cell cycle expression system. Using the PEO1 FUCCI cells to assess performance, we achieved an accuracy of 89% using our simple classifier on a balanced data set of 288 cells each in G1 and S phase, respectively, where we use 200 cells in G1 phase to obtain cutoff thresholds, and used these thresholds to evaluate the prediction on 85 cells each in G1 and S phase (Fig. 3). We select the G1 cells in order to make the analysis comparable to the PERT approach, and because the data set is highly enriched for S phase cells. Our results compared favourably to PERT, a recently released updated cell cycle prediction method [60], where we achieved 71% accuracy on the same data with the early release version of the software.

**Fig. 3:**
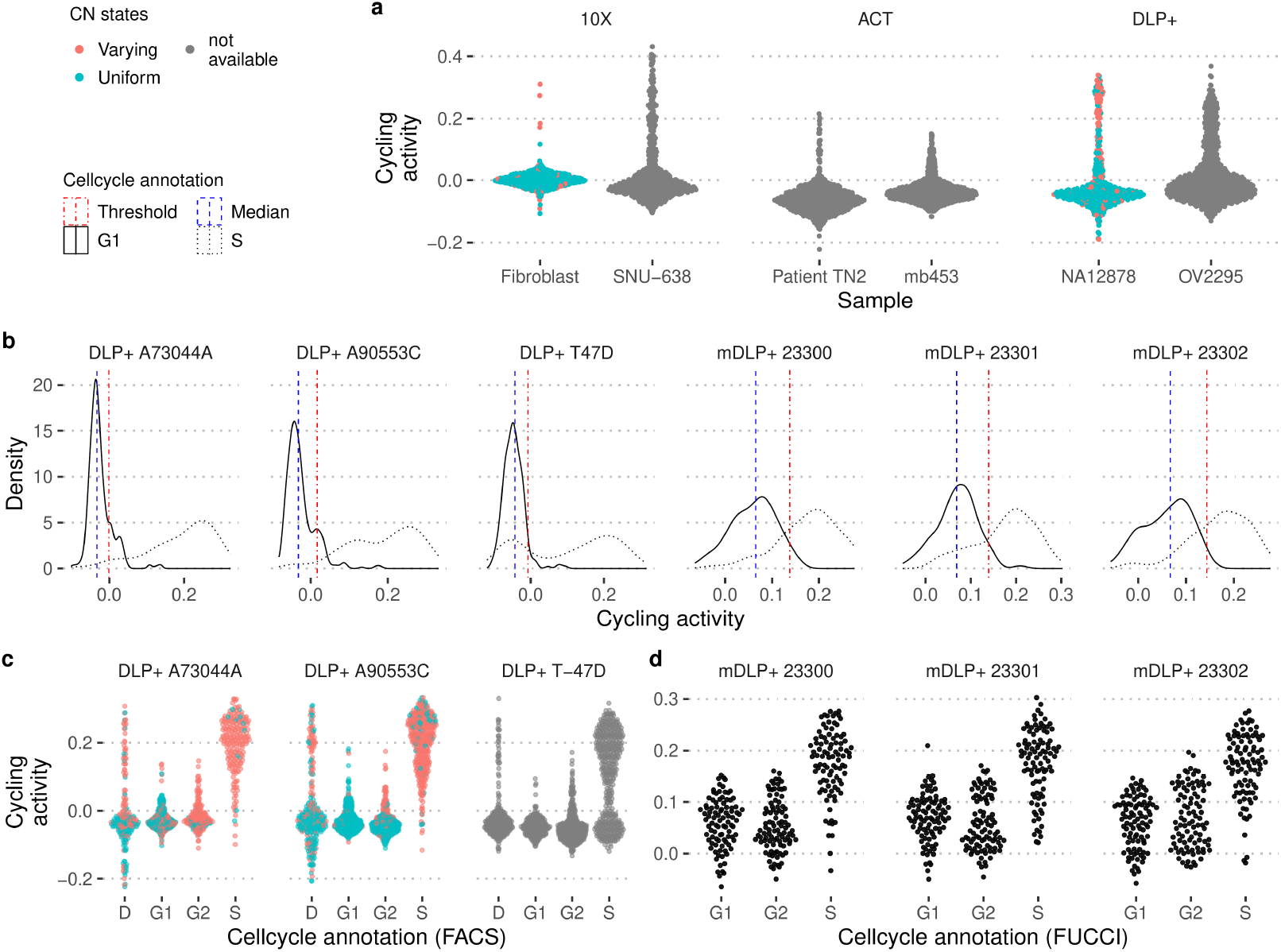
Detection of replicating cells across different sequencing technologies and cell lines. **a)** Distribution of cycling activity in samples sequenced with three different sequencing technologies: 10X, ACT and DLP+. Uniform and varying CN state indicates whether the copy number calls indicate a mostly diploid genome (as in G1/G2), or cycling cells (non-uniform CN state), respectively. **b)** Distribution of cycling activity for G1 (solid line) and S phase cells (dotted line). The median of the distribution (blue line) is estimated and the left side of the distribution is used to determine a standard deviation that covers the majority of cells in G1 phase (red lines) and to determine an appropriate cutoff value to identify cells in S phase. **c)** Cycling activity for normal, diploid cells (A73044A and A90553C), and a cancer cell line (T-47D) sequenced with DLP+ technology. The cell cycle annotation is based on sorting cells with FACS using DAPI staining and grouping cells either into apoptosis state (D), or in the three cell cycle stages (G1, S phase, G2). This measure can only be reliable estimated for normal, diploid cells, and is not applicable in other samples. **d)** Cycling activity for PEO1 FUCCI cells that have been sorted into different cell cycle stages based on the FUCCI marker system. The cells have been sequenced with a modified version of the DLP+ protocol (mDLP+).

In order to further validate the approach, we examine two other examples. First, we considered normal diploid cells (10X Fibroblast and DLP+ NA12878). In this case, we could assume that the vast majority of CNAs were due to cells undergoing DNA replication. We observed that the majority of normal cells with many CNAs (highly indicative of cells undergoing S phase) scored high on the cycling activity measure (Fig. 3). Similarly, across cell lines and libraries, we observed the typical asymmetric distribution of cycling activity. The only exception (10X Fibroblast cells) were cell cycle arrested, and so we did not expect to observe any cycling cells in this set.

Looking at a further set of 10 cell lines sequenced with the 10X technology we observed a robust negative relationship between doubling time of a cell line and our estimate of proportion of cycling cells (r = -0.75, p = 0.02, see Fig. S7).

### Benchmarking initial ploidy estimation

A major limitation of using all four steps of *scAbsolute* is that it requires data produced without whole-genome amplification at sufficient read depth. However, this requirement only affects step 4 and the initial ploidy estimation in the first three steps of *scAbsolute* can be applied more widely.

Here we evaluated *scAbsolute*’s initial ploidy estimation (steps 1-3) on 3 published datasets where orthogonal information was available for the ploidy state. These data covered a wide range of realistic copy number profiles and technologies (Fig. 4). The first dataset, consisted of cell lines profiled using the 10X single cell DNA protocol, where we had karyotype estimates available [59]. The second dataset consisted of cell lines profiled using DLP+ [28], with the known ploidy estimates of the cell lines from the literature. The final dataset profiled using the ACT protocol [34] had cells selected using DAPI staining and FACS. The advantage of the ACT data was that the ploidy estimates covered the entire population, making it possible to compare different methods and ascribe ploidy outliers to wrong ploidy estimates rather than cell-level variation.

**Fig. 4:**
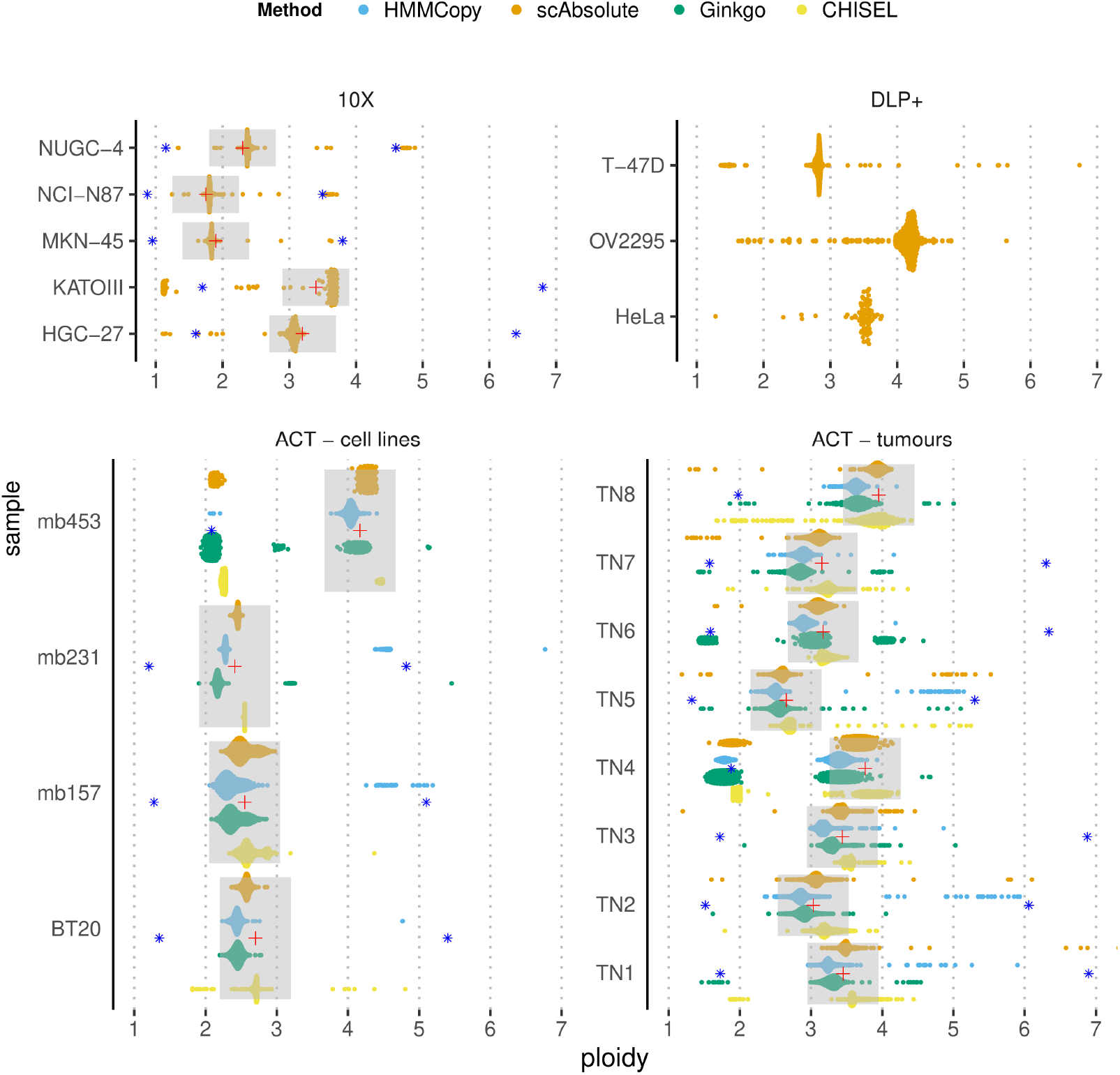
Ploidy prediction results for 4 methods (*scAbsolute*, HMMCopy, Ginkgo, and CHISEL) across single-cell DNA sequencing datasets from 3 different sequencing technologies (10X, DLP+, ACT) from cell lines and tumour samples. We only ran steps 1-3 of the *scAbsolute* algorithm. The data sets used are described in detail in the Methods section. Experimental ploidy annotation for the 10X data was based on karyotype information, and for the ACT data was based on FACS sorted cells using DAPI staining. Grey ranges of ±0.5 around the experimental point estimate (red cross) of the sample ploidy are indicated. Blue asterisks indicate ploidy levels of 1*/*2 or 2 times the experimental ploidy estimate. For DLP+ data, no experimental ploidy annotation was available, but estimates were in accordance with ploidy estimates for the respective cell lines based on public karyotype information.

We assessed performance in two ways: first, as the percentage of cells outside an experimental ploidy window of ±0.5 around the peak of the DAPI distribution (for the ACT dataset). This window included uncertainty from segmentation and FACS sorting, but excluded true ploidy changes. Second, as the mean absolute distance across all cells in a sample from the experimental ploidy estimate. Table 1 gives a detailed overview over the prediction results. For both metrics, we found that scAbsolute consistently predicted the correct ploidy solution for the large majority of cells.

**Table 1:**
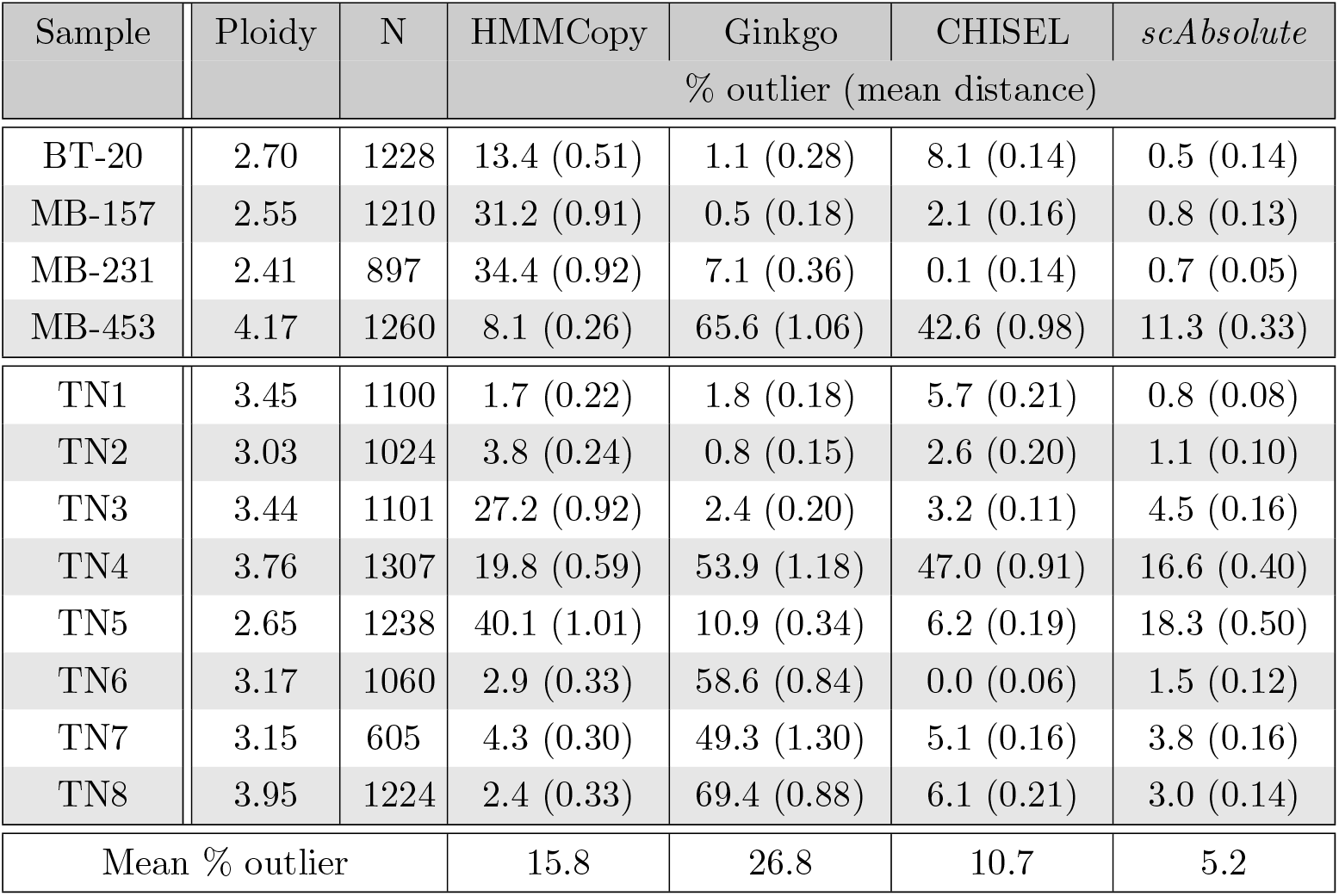
Comparison of ploidy prediction for ACT samples. Ploidy prediction based on *scAbsolute*, Ginkgo, HMMCopy, and CHISEL. Ploidy estimate is based on FACS sorting cells using DAPI staining. We considered an estimate to be an outlier if it fell outside of a ±0.5 window around the experimental ploidy point estimate. We also gave the mean of the absolute distance in ploidy between the experimental point estimate and the computational estimate across all cells in parenthesis. The data were all sequenced with the same technology (ACT) and published in [34]. N indicates the number of cells per sample. The last row gives the mean % of outliers across all samples per method.

We also used the ACT data to compare performance of different computational tools, assuming that ploidy outliers are indeed due to erroneous ploidy prediction. We compared scAbsolute to Ginkgo [55], HMMCopy [50] and CHISEL [58] (Fig. 4). Unlike the other methods, CHISEL is using information from SNPs to improve ploidy calling. In our case, we only had access to normal reference samples in the form of WGS bulk-exome sequencing for the ACT tumour samples (TN1-8), and this data was insufficient to call variants for the normal CHISEL workflow. However, we were able to run CHISEL in a nonormal mode, where SNPs were called based on the pseudobulk generated from the single cells and filtered and phased via a reference database (here we use TOPMED [63]), and we used this mode in our analysis of the ACT data. On average, scAbsolute was wrong in 5% of cases, CHISEL had wrong ploidy predictions in 11% of cases, HMMCopy in 16%, and Ginkgo in 27%. In only one case (TN5), scAbsolute performed considerably worse than Ginkgo with an error rate of 18% compared to Ginkgo’s 10%. This case, and a similar scenario in the TN4 sample, were due to the unidentifiability of the problem, that is not addressed in steps 1-3 of *scAbsolute*. Note that the ACT read data were based on single-read sequencing and given this fact, they did not have sufficient coverage to apply step 4 of *scAbsolute*. The average above hides considerably variation across samples. The three samples with the highest percentage of wrongly assigned ploidies were 40% and 34%, and 31% in the case of HMMCopy (TN5, MB-231, and MB-157), 69%, 66%, and 59% for Ginkgo (TN8, MB-453, and TN6), and 47%, 43% and 8% (TN4, MB-453, BT-20) for CHISEL, compared with 18%, (TN5) 17% (TN4), and 11% (MB-453) for *scAbsolute*.

CHISEL performed best among the methods we compared to, with a substantial number of outliers in only two samples. In addition, we ran our algorithm on one of the samples originally published with CHISEL that includes raw sequencing data for about 10 000 cells. Here, we could run CHISEL in its original implementation, without recourse to the *nonormal* mode. In general, we found good overlap between the two predictions, in the form of identical predictions (predictions on the diagonal, see Fig. S8). For this sample, we did not have experimental ploidy estimates available, however, we conducted a comparison of outliers to identify differences between the methods. First, we randomly selected cells that were predicted to be nearly diploid by *scAbsolute*, and had a ploidy predicted to be greater than 3 by CHISEL. We observed CHISEL selected higher ploidy solutions, that were not necessarily supported by copy number levels in some of these instances (Fig. S9), whereas *scAbsolute* seemed to fail in some cases where the underlying segmentation was wrong. Second, we compared cells that were predicted to be nearly diploid by CHISEL, and had a ploidy predicted to be greater than 3 by *scAbsolute* (see Fig. S10). Here, we observed a few cases of highly uneven cell coverage (possibly indicating failed sequencing runs) and some failures of *scAbsolute* to detect the correct solution in very uniform samples (the rate of these cases is very low at about 1%). We also compared the overall copy number profiles for section E of patient S0, as presented in Zaccaria and Raphael [58] and found overall similar copy number predictions in this particular case (Fig. S11).

## Discussion

The increasing availability of low-coverage whole-genome sequencing of thousands of individual cells offers an opportunity to study tumour heterogeneity and evolution at an unprecedented resolution. *scAbsolute* is a novel tool to investigate cell ploidy and replication status at the single-cell level. We demonstrate the general applicability of *scAbsolute* across three recently released single-cell DNA sequencing protocols, and show a practical advantage of using whole-genome amplification free protocols (DLP+, ACT) in detecting WGD events. In addition, *scAbsolute* has potential use in the problem of doublet detection in future protocols that combine a droplet-based single-cell isolation with whole-genome amplification free library preparation.

The approach is fundamentally different from other approaches originally developed in bulk sequencing settings and recently extended to the single-cell domain [58] that use B-allele frequency (BAF) in order to help identify ploidy solutions. However, by using cell specific haplotype counts, and the high quality total copy number predictions provided by *scAbsolute*, it is possible to estimate allele and haplotype specific copy number states, as recently demonstrated [52, 58]. In particular, it is straightforward to estimate the allele specific copy numbers using the BAF as estimator, once the total copy number is known.

One limitation of the computational approach is the inability to distinguish cases of cells in G2 phase of the cell cycle from WGD cells. However, it is possible to address this either via recourse to estimates of the relative number of cells in G2 state, or via integration with experimental evidence, such as Geminin staining. Similarly, the identification of WGD/G2 cells is possible only in cases of paired-ends reads, with sufficient read depth and using whole-genome amplification free sequencing protocols. It might potentially be possible to extend this approach to single-end reads. However, we believe that this would require a substantial increase in read depth and there is currently no public data available to test this hypothesis.

The identification of per-cell ploidy and with it per-cell read depth lies at the basis of many downstream applications, such as copy-number segmentation, estimation of allele- and haplotype-specific copy number states, and the building of tumour phylogenies based on copy-number profiles. By solving the ploidy problem, further progress in developing more accurate and reliable CNA calling methods seems feasible.

We offer an alternative to the use of FACS with DAPI staining to identify cell populations at different ploidy levels. This removes the additional cost and time involved in the FACS step and opens the way for a more unbiased investigation of intratumour heterogeneity. In particular, the choice of cutoffs for FACS might lead to a bias in selecting more homogeneous cell populations and lead to an underestimation of the true level of heterogeneity in tumours. FACS might still be useful in order to reduce the amount of non-informative normal cells sequenced, thus reducing overall costs, and as an experimental validation, however.

In addition to the problem of ploidy, we also address the issue of replicating cells. Here, we present a simple, but robust measure of cycling activity and we demonstrate its applicability across a range of cell lines and sequencing technologies. Importantly, we demonstrate some challenges of using DAPI staining based measures of cell cycle status and how this can bias the training of cell cycle classifiers.

Another important aspect of single-cell methods is their applicability to heterogeneous samples, such as observed in complex tumours. We have developed a tool that is robust across different samples and sequencing protocols. In the case of highly heterogeneous samples, it is also possible to conduct a clonal decomposition based on copy number profiles prior to subsequent detection of ploidy and replication status. In particular, we believe this to be favourable to selection of subpopulations based on FACS sorting, as this can introduce a bias in the representation of genomic heterogeneity. However, more work will be required to systematically evaluate the best techniques to study intratumour heterogeneity at the single-cell level.

*scAbsolute* is the first computational approach to overcome the unidentifiability problem associated with WGD events in copy number calling of single cells. This is particularly relevant for the study of early tumourigenesis, by making it possible to study WGD and other ploidy changes in small lesions with very limited numbers of cells. *scAbsolute* offers an unbiased way of estimating tumour ploidy and identifying replicating cells; this will help to further improve the quality and reliability of copy number calling in single-cell DNA sequencing data. It is an important contribution to the unbiased measurement of intratumour heterogeneity. Ultimately, it is a further step towards elucidating the role of CIN in cancer.

## Methods

### Bin-level quality control

Prior to any analysis, we create a set of high quality genomic regions extending existing bin-annotations and specifically targeted at single-cell DNA-sequencing (scDNA-seq) data. We use these novel bin annotations across all further experiments and they are available as part of the scAbsolute software. The primary purpose of this step is to only include genomic bins for which the assumption of a linear relationship between copy number status and observed read counts holds.

The set is created by combining data from three different sequencing technologies (10X, DLP+, and an in-house protocol referred to as JBL), comprising more than 7000 diploid cells (We do not have access to diploid cells sequenced with the ACT protocol). In order to create a single, unified set of bin annotations, we conduct below analysis separately for the three sequencing technologies, and exclude all bins that fail the quality criterion in any single sequencing technology (see Fig. S12).

To determine bin quality, we only look at diploid cells, so we can disregard the issue of copy number status and segmentation. The set contains no cells in S phase or in G2/M phase (based on FACS sorting, and cell cycle arrest in the case of the 10X cells). In order to have sufficient reads per bin to reliably detect any deviations, and at the same time to have as high as possible a genomic resolution, we conduct this analysis on reads binned at 100 kb resolution.

First, we normalise the per-cell reads by dividing the number of reads in each bin by the expected number of reads per bin across a cell, creating a normalised read per bin value. Subsequently, we use the median of the normalised reads per bin across all cells sequenced with the same sequencing technology as a per-bin quality metric. Initially, we remove all bins that have a median value of more than 4 or less than 0.10 per bin, and a mappability value smaller than 70 (11% of bins). Note, that the expected value would be 1, so a value of 4 corresponds to a median number of reads falling into a bin four times higher than would be expected on average.

In a second stage, we look at the relationship between GC content and mappability, and median normalised read counts for each of the sequencing technologies. Separately for each technology, we fit a Generalised Additive Model to characterise the relationship between GC content and normalised read counts, and mappability and normalised read counts, and remove all data points that deviate more than 2 standard deviations (see Fig. S12). In addition, we use a kernel density estimate to select regions of high density, and remove all cells outside the high-density regions. By using these two criteria, we aim to select only high quality bins that have minimal read count deviation that is not explained by GC content and mappability. Lastly, we use maps of centromere and telomere regions to specifically filter parts of these regions that have not been filtered in the previous steps.

Overall, we remove about 16% of the bins, containing about 9% percent of total reads on the autosomes. For the X and Y chromosomes, we use the existing QDNAseq annotations and run a simplified version of the above pipeline to remove outliers on the X chromosome (only based on density estimates). In the case of the Y chromosome, we remove bins solely based on deviations in the total number of reads observed, because of the low number of reads on these chromosomes in our data (Fig. S13).

### Initial Segmentation with unknown ploidy

We use a dynamic programming approach based on the PELT algorithm [61] to find an initial segmentation of our read counts. We model the read counts with a Negative Binomial distribution [64].

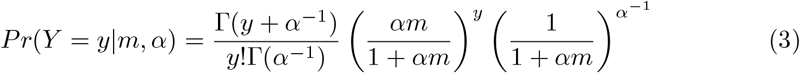

Here, *α* denotes a measure of increasing overdispersion; for values of *α*→ 0, the distribution converges to that for the Poisson [65].

There exists no analytical solution to the Negative Binomial Maximum Likelihood estimate of the overdispersion parameter. We therefore choose a Methods-of-Moments estimator for the parameter *α* in the cost function [66]. We use this initial segmentation as a stepping-stone in identifying a cell’s correct ploidy, and are not interested in an optimal segmentation at this early stage, but rather a robust and reasonably fast approach that achieves reasonable accuracy. We note that optimal segmentation is not the focus of this manuscript, and it is possible to replace the segmentation algorithm with one of the user’s choice.

The cost function for a segment *y*_*j*_, …, *y*_*j*+*k*_ of length k is defined as

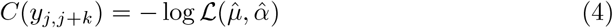

where 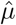 and 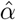 are the parameters of the Negative binomial likelihood estimated from the data and ℒ denotes the negative binomial likelihood function. The total cost is then the sum of the segments.

The mean is estimated via 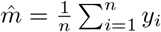, and the overdispersion is estimated as

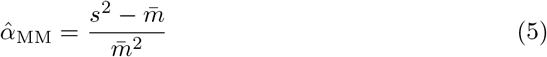

where 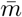 and *s*^2^ are defined as sample mean and variance, respectively [66].

### Ploidy estimation

Ploidy estimation describes the identification of copy number segments with underlying copy number states. This is equivalent to finding a scaling factor *ρ*, so that we can directly measure the ploidy *p* of a cell.

Note that we assume that observed reads scale linearly with copy number state, e.g. we expect to observe on average twice as many reads for copy number state 2 than for copy number state 1, and so on. In our approach, given an initial – not-necessarily exact – segmentation, we consider the marginal distribution of the segmented copy number states, i.e. the spatial correlation of neighbouring bins has been accounted for through the initial segmentation. Since we are dealing with individual cells, in theory all copy number states should appear at integer values (with the exception of bins that contain two or more different copy number states), with some states possibly not observed in a cell (see Fig. 1 for an example for the marginal distribution of segmented counts). In a normal diploid cell which has no CNAs, we would expect to observe a single value scaled by a scalar value 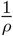 denoting the read depth of the cell (excluding the sex chromosomes). However, given the inherent measurement noise and imperfect segmentation, in fact one observes a distribution of values around the integer states.

Here, we assume the errors are normally distributed and approximate the marginal distribution of the segmented counts with a (constrained) Gaussian Mixture Model. The constraint on the Gaussians is based on the fact that instead of estimating *K* means *μ*_1_, …, *μ*_*K*_, we restrict the location of the means to *μ*_1_ = 1 *· ξ*, …, *μ*_*K*_ = *K · ξ*. Consequently, we estimate a single parameter *ξ* and a set of *K* standard deviations *σ*_*k*_ for the *K* Gaussians in the model. We might not observe all possible states and we don’t know how many states we will observe in any given cell. Consequently, we model the appearance of individual clusters with a Dirichlet Process. The variational distribution of the Dirichlet Process is truncated at *T* components [67]. In order to further speed up the computation, we use stochastic variational inference [68] and implement the algorithm in TensorFlow [69]. Overall, we estimate the posterior probability 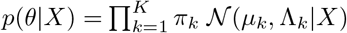. Details and the mathematical derivation of the model updates can be found in the appendix.

A major challenge with estimating absolute copy number states is the unidentifiability of a unique solution. There exist many potential solutions for each copy number profile, as it is always possible to shift or scale the solution. For example, consider the case of a perfectly diploid and tetraploid genome. Both are biologically plausible, e.g. in case of a Fibroblast cell in G1/G2 phase, but indistinguishable without any additional mathematical constraints. From a mathematical point, even a biologically implausible triploid solution is equally possible. This problem is less relevant in cancers with a high number of CNAs, as one tends to observe many of the copy number states in a single cell, thus making it easier to identify the correct solution. This makes the problem considerably easier, however, it would still not be possible to detect evolutionary recent whole-genome doubling events or distinguish between cells in G1/G2 phase. As a consequence, the design of our model does not enforce any particular solution. Instead it returns one possible solution, with the constraint, that the means of the Gaussians, i.e. the observed copy number states, occur at a distance that is an integer-multiple of an arbitrary unit distance and the set of solutions lies within a biologically reasonable, user defined ploidy range.

In order to select a biological plausible solution *ρ* out of the discrete set of possible values 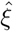 within the given ploidy range, by default, we select the solution that has a minimal least squared error. In practice, this tends to be the solution with the lowest ploidy within the given ploidy range. For a biologically plausible ploidy range, we chose a minimum ploidy of 1.1. The assumption here is that cells with more copy number loses are probably not viable. As an upper bound, we chose a ploidy of 8 by default. However, this can be flexibly adjusted given other sources of information.

### Addressing the unidentifiability problem

Up to this point, the approach cannot distinguish between cells in G1 and G2 phase of the cell cycle, or cells directly after WGD. Here, we demonstrate that we can in fact reliably differentiate between these cells, given we have a i) sufficiently high read coverage ii) we use a sequencing technology without a whole-genome amplification step and iii) paired-end read sequencing. Read coverage depends both on the ploidy of the sample, and on the size of the genomic regions that is covered (and mappable to) sequencing reads. It might be possible to avoid the need for paired-end read sequencing, if the coverage is substantially higher. However, we do not have access to any data to test this scenario, and it would be relatively expensive to acquire it given the depth required.

The basic idea behind the approach is to analyse the number of overlapping reads across the genome. For this purpose, we compute a measure of how densely reads are located on the genome, and how many reads physically overlap. Because the genome is not amplified, we then can assume that the number of overlapping, unique reads is directly proportional to and limited by the number of physical copies of the genome at the given location (Fig. S2). The only other two parameters we expect to influence the observed read density are the cell ploidy and the per-cell read depth (as measured by the *ρ* value).

Initially, for each properly mapped read, we count the number of overlapping reads across the genome, using the start and end position of paired-end reads as genomic start and end position, respectively. The size of the regions used is then determined by the fragment size. We compute the average number of overlapping reads across genomic bins, referring to it as read density. In order to account for varying copy number status, to make the estimation process more robust and to enable the comparison across cells with different copy number profiles, we fit a robust linear regression model to approximate the relationship between copy number state and read density (Fig. 1). We also split the data by chromosome, weighting each data point by the number of the bins covered. In order to directly compare cells, we use the predicted value of read density for a copy number state of 2 as per-cell measure of read density. Note that we can even predict this value in the absence of any observed copy number state 2. In the case of normal, diploid cells, we resolve to using the median read density, as the linear regression would otherwise be underdetermined with only a single data point at a copy number state of 2.

In a second step, we use a reference data set of cells with known ploidy to create a model of expected read density for a given read depth. Here, we combine cells with known ploidy and sequenced with either the DLP+ or the mDLP+ technology in order to create a reference model of expected read density for a given value of *ρ* (Figs. 2a and 2b). We can then compare the residual read density, measured as the deviation between observed and expected read density at a given value of *ρ* in order to distinguish cells that have been assigned a wrong ploidy (see Figs. 2c and 2d).

### Detection of replicating cells

We devise a simple test statistic to examine if a cell is in the S phase of the cell cycle. The test is based on differences in replication timing for different parts of the genome. In order to quantify replication timing per genomic bin, we use an existing annotation from the Repli-chip from ENCODE/FSU project [70, 71], and average the replication times determined in these experiments across multiple cell lines and across genomic regions contained in a bin. This allows us to obtain a single measure of replication time per genomic bin.

It is known that GC-content and mappability lead to a bias in the read counts observed across different genomic locations [55, 72]. Here, we fit a Generalised Additive Model (GAM) to estimate and correct for the bias individually per cell. The advantage of this approach is that it gives us a direct estimate for the impact of the covariates on the mean of the read counts observed in each bin. We model the observed read counts x for every bin *j* as *x*_*j*_ ∼NegBin(*μ*_*j*_, *α*), with *log*(*μ*_*j*_) = *β*_0_ + *β*_1_*ν* + *s*(*gc*_*j*_, *map*_*j*_), where *ν* denotes the segmentation value of a bin, i.e. the median value of read counts across multiple neighbouring bins and *gc*_*j*_ and *map*_*j*_ the GC content and mappability values in bin *j*, respectively. We model the impact of varying GC and mappability content jointly with thin plate regression splines. We obtain coefficients of the impact of GC content and mappability variation on the mean expression for every cell and every bin, in the form of the coefficients *s*(*gc*_*j*_, *map*_*j*_) that directly relate to the mean-expression and we use these values as covariates below.

We perform a partial correlation trend test using the *Spearman* rank correlation statistic [62, p. 882], computed for each segment *S*_*l*_ within a single-cell copy number profile. Each segment is a series of consecutive bins with identical copy number state, as identified by the segmentation algorithm. The trend test is performed on the raw count data within a given segment, sorted by increasing replication time, while controlling for the per bin GC-content and mappability value via the coefficient *s*(*gc*_*j*_, *map*_*j*_) described above. We refer to the value of the test statistic for each segment *l* as *τ*_*l*_. Subsequently, the median over all the partial correlation test statistics *τ*_*l*_ weighted by the respective segment length is computed. We refer to this single measure as cycling activity. In general, we observe positive values as indicative of cells in S phase, undergoing replication. The distribution of the cycling activity measure is symmetric around its mode with an additional long tail, that we identify with the replicating cells (see Fig. 3).

We make use of the characteristic shape of the distribution to classify a cell’s replication status across different sequencing technologies and libraries. We chose the threshold dynamically by identifying the median of the distribution and determining the standard deviation using only the left (negative) part of the cycling activity distribution. By default, we use a cutoff corresponding to 1.5 standard deviations from the median of the distribution as the threshold for classifying a cell as being in S phase (Fig. 3b). To detect additional, individual outlier cells, we compute a global measure of mutual information between observed reads and replication time per bin, conditioned on the copy number state of a bin. We use an entropy estimator based on the empirical entropy distribution. We exclude outlier cells that are more than 1.5 times the interquartile range away from the third quartile of the mutual information measure (boxplot criteria for individual outliers). This makes the approach easily applicable to new single-cell data without cell cycle annotation, and without having to adapt the cell cycle classifier to a new feature distribution in each instance.

## Experimental data

### Cell culture

The PEO1 cell line, derived from an ovarian adenocarcinoma [73, 74], was cultured in RPMI-1640 + 10% heat inactivated fetal bovine serum + 2 mM Sodium Pyruvate (Invitrogen) at 37 ^*°*^C, 5% CO2. The PEO1 cell line was a gift from Simon Langdon. The PEO1-STOP cell line is a PEO1 clonal derivative containing a hemizygous BRCA2 stop codon mutation BRCA25193C/G (Y1655X) [74, 75]. PEO1-STOP cells were cultured in RPMI-1640 + 10% heat inactivated fetal bovine serum (Invitrogen) at 37 ^*°*^C, 5% CO2. Cell lines were routinely screened by Phoenix Dx Mycoplasma Mix qPCR (Procomcure Biotech), and tested mycoplasma negative. Cell line identities were authenticated by STR genotyping.

### Preparation of PEO1 STOP-FUCCI cell line

Fluorescent ubiquitination cell cycle indicator (FUCCI) [76] is a lentiviral expression system that can be used to identify cell cycle status by fluorescently tagged Geminin, expressed in S and G2 phase, and Cdt1, expressed in G1 and early S phase. To generate a experimental tool to validate *scAbsolute’s* ability to detect cells in G1 vs G2, the all in one FastFUCCI lentiviral expression system [77] was transduced into PEO1-STOP cells to generate cells expressing Cdt1-mKO2 and Geminin mAG [78]. The FastFUCCI cassette contains a puromycin selection marker to allow for positive selection of FUCCI expressing cells.

To generate the FastFUCCI lentiviral particles, the pBOB-EF1-FastFUCCI-Puro cassette was transfected into 293FT cells with pVSV-G, pRSDV, and pMDLg/pRRE as previously described [77]. Viral Supernatant was collected and concentrated by centrifugation at 3000 g for 30 minutes at 4 ^*°*^C using an Ultra-15 centrifugal filter (Amicon). To generate the PEO1-STOP cells expressing FUCCI, PEO1-STOP cells were seeded at a concentration of 5 *×*10^4^ cells per well of a 12-well plate and then incubated overnight at 37 ^*°*^C with 250 μL concentrated viral supernatant and 10 μg*/*μL polybrene. Following this incubation, the viral supernatant was removed, and replaced with fresh media. The cells were assessed visually 72 hours later for fluorescent protein expression prior to positive selection with 0.5 μg*/*mL Puromycin for 96 hours.

### Flow Cytometry validation of PEO1-STOP FUCCI

To validate the expression of Geminin-mAG and Cdt1-mKO2, the cells were assessed by Flow Cytometry using the FACSMelody cell sorter (BD Biosciences). Side scatter area (SSC-A) versus forward scatter area (FSC-A) was used to gate cells from debris. Side scatter height (SSC-H) versus side scatter width (SSC-W), and forward scatter height (FSC-H) versus forward scatter width (FSC-W) was used to gate singlets from doublets.

To assess the cell cycle fluorescence, mAG and mKO2 staining was compared to cytometric cell cycle analysis using DAPI staining. DAPI intensity directly correlates to DNA content, so can be used to identify G1, S and G2 cells [79]. PEO1-STOP FUCCI cells were prepared as below for DAPI staining. DAPI vs mKO2 shows high expression of mKO2 in DAPI gated G1, correlating to high Cdt1 expression, with low expression of mKO2 in DAPI gated S and G2. DAPI vs mKO2 shows high expression of mAG in DAPI gated S and G2, correlating to high Geminin expression, with low expression of mAG in DAPI gated S and G2 (see Fig. S6).

### Cell cycle sorting using DAPI, Geminin, Cdt1

Fresh cell suspension was pelleted and resuspended in 1 mL PBS per 5 million cells. 4 mL ice cold, 100% Methanol was added per million cells drop-wise to the cell suspension whilst agitating the cells, then incubated for 15 minutes at −20 ^*°*^C. Following fixation, the methanol was removed from the cell pellet, and the cells were resuspended in 1.25 mL ice cold PBS per million cells. Permeabilisation and blocking of the cells occurred via a 15 minute incubation with 1 mL PBS supplemented with 10% Normal Goat Serum (Abcam) and 0.1% Triton-X per million cells.

For Geminin Antibody staining, cells were incubated overnight at 4 ^*°*^C, 300RPM in Primary Antibody Solution (Geminin E5Q9S XP Rabbit mAb (#52508, Cell Signalling Technologies)) diluted 1:100 in PBS plus 10% Normal Goat Serum) at a ratio of 100 μL per 1 million cells. Cells were then incubated for 1 hour at 22 ^*°*^C, 300RPM in a 1:500 dilution of Alexa Fluor 488 secondary antibody (Goat anti-Rabbit IgG (H+L) Cross-Adsorbed Secondary Antibody, Alexa Fluor™488 #A-11008, Invitrogen). For DAPI staining, both PEO1 and PEO1-STOP FUCCI were incubated in 4 μg*/*mL DAPI solution (1 mg*/*mL, Thermo Scientific) in PBS for 10 mins 1 mL per 1 million cells.

PEO1-STOP FUCCI cells were sorted into G1, S, and G2 populations using FACSMelody cell sorter (BD Biosciences) into 1.5 mL protein Lo-bind tubes (Eppendorf). Cell cycle gating was performed as above using a combination of DAPI staining, measuring DNA content, and FUCCI status. G1 cells were gated using DAPI staining and mKO2 positive staining. S and G2 cells were gated using DAPI staining and mAG positive staining.

For PEO1 multiple cells assay, PEO1 singlets were gated as above, and further gated for G1 status using a combination of DAPI and Geminin antibody staining. Geminin positive and negative populations were sorted for use as controls for cellenONE (Cellenion) gating.

### Single-cell isolation using cellenONE

Single-cell sorting was performed using an image based piezo-acoustic nanolitre dispenser (cellenONE, Cellenion). Single cells were selected based on cell line specific parameters of cell diameter, elongation and circularity (see Table S2). For PEO1 ploidy experiments, active selection of cells in G1 cell cycle state was based on positive selection of DAPI (blue channel), and negative selection of Geminin (green channel). For FUCCI experiments, flow sorted populations of G1, S and G2 were dispensed by the cellenONE using positive selection of DAPI only (blue channel). Cells were dispensed into 384 well plates containing pre-dispensed 300 nL of 30 mm Tris buffer, then stored at −20 ^*°*^C.

### Single-cell shallow whole-genome library preparation

Single cell whole genome direct library preparation was prepared based on DLP+ [28] with the following modifications (mDLP+). Plates with dispensed cells were removed from the freezer and proceeded to cell lysis by adding freshly prepared 300 nL of 2*x* lysis buffer (30 mm Tris pH8.0, 1% Tween-20, 1% Triton X-100) to each well using Mantis Liquid Dispenser (Formulatrix) followed by 10 minutes incubation at 55 ^*°*^C, 15 minutes at 75 ^*°*^C. DNA Tagmentation was performed by adding 1 μL of TD buffer and μL of ATM buffer mixture (Nextera XT DNA Library Preparation Kit, Illumina) and incubated at 55 ^*°*^C for 5 minutes. The tagmentation reaction was neutralised by 5 minute incubation at room temperature with 0.5 μL NT buffer (Nextera XT DNA Library Preparation Kit, Illumina). Library amplification was performed by adding 4 μL (2X) Equinox Amplification Master Mix (Equinox Library Amplification Kit, Watchmaker Genomics) and unique dual indexed adapter pool of 2 μm (IDT for Illumina DNA/RNA UD Indexes, Illumina), using following PCR conditions: 72 ^*°*^C for 5 min, 98 ^*°*^C for 45s, 12 cycles of 98 ^*°*^C for 15s, 60 ^*°*^C for 30s, 72 ^*°*^C for 45s, followed by 72 ^*°*^C for 5 min. Lid temperature 105 ^*°*^C.

### Single-cell shallow whole-genome library pooling, normalisation and sequencing

#### PEO1 ploidy experiment

To assess the accuracy of *scAbsolute* we created experimental models reflecting different ploidy status by combining different numbers of cells, for example: 3N (1 cell), 6N (2 cells), etc. The single cell library prep was performed as above, libraries from the same number of input cells were pooled together and purified using 1X AMPure XP (Beckman Coulter) beads. Sample transfers were performed by mosquito HV (SPT Labtech). The purified libraries were quantified by Qubit dsDNA HS (Invitrogen) and pooled in equimolar ratios. The final library pools were purified again with 1X AMPure XP (Beckman Coulter) beads, evaluated by Tapestation (D5000 ScreenTape Assay) and quantified using KAPA SYBR FAST universal kit (Roche) prior sequencing. Final libraries were sequenced using PE-50 mode on NovaSeq6000 (Illumina) instrument aiming for 6 million reads per cell.

#### PEO1-STOP FUCCI

Single Cell libraries for PEO1-STOP FUCCI were performed as above, pooled after the library amplification and purified using 1X AMPure XP (Beckman Coulter) beads, evaluated by Tapestation (D5000 ScreenTape Assay) and quantified using KAPA SYBR FAST universal kit (Roche) prior sequencing. Final libraries were sequenced using PE-50 mode on NovaSeq6000 (Illumina) instrument aiming for 6 million reads per cell.

### Pooling and Normalisation

For PEO1 ploidy experiments investigating increasing number of cells inputted for mDLP+, pooling of cells for sequencing was determined by both volume and mass. Initially libraries were pooled into individual groups dependent on number of cells inputted into the reaction A 1X Ampure bead clean-up (Beckman Coulter) was performed, and then the pools were quantified by Qubit HS (Invitrogen). The pools were then normalised by mass of DNA multiplied by the number of cells inputted per mDLP+ reaction to normalise number of sequenced reads/cell rather than number of reads/reaction. The normalised pool was then subjected to a final 1X Ampure bead clean up prior to sequencing. For PEO1-STOP FUCCI experiments cells were pooled by volume prior to two 1X Ampure Bead purifications.

### Sequencing

Analysis of pooled, purified single-cell libraries was performed using either Tapestation DS5000 ScreenTape Assay for Tapestation (Agilent) or DNA High Sensitivity Kit for the Bioanalyzer 2100 (Agilent). Libraries were sequenced on the Illumina NovaSeq (paired-end 50bp reads) by the Genomics Core Facility at CRUK Cambridge Institute, University of Cambridge.

## Declarations

### Ethics approval and consent to participate

Not applicable

### Consent for publication

Not applicable

### Availability of data and materials

The PEO1 FUCCI and artificial multiplet sequencing data (3N-15N) generated as part of the study are available in the European Nucleotide Archive under accession number PRJEB61928.

The other sequencing data used cover three different single-cell sequencing technologies: 10X data [59] is available in the NCBI Sequence Read Archive; accession number PRJNA498809 (cell lines NUGC-4, NCI-N87, MKN-45, KATOIII, and HGC-27; these cell lines were selected because they had sufficient read depth for our analysis), and two demonstration datasets (a normal diploid Fibroblast cell line, and a Breast Tumour Tissue sample) published online as part of the 10X Single Cell DNA sequencing technology demonstration (https://www.10xgenomics.com/); DLP+ data [28] is available in the European Genome-Phenome Archive under accession number EGAS00001003190 (see Table S1 for an overview) and ACT data [34] is available in the NCBI Sequence Read Archive under accession number PRJNA629885 (cell lines BT-20, MB-157, MB-231, MB-453, and tumour samples TN1-8). We excluded samples for which the majority of data is below the 25 *ρ* threshold at 500 kb resolution to restrict the influence of segmentation on the ploidy calling.

### Availability of code

The source code for *scAbsolute*, and scripts to reproduce all figures and analyses, and to reproduce results for other tools is available at https://github.com/markowetzlab/scAbsolute. We also provide a snakemake workflow and tutorial to run the tool at https://github.com/markowetzlab/scDNAseq-workflow. *scAbsolute* uses the package environment and genome annotations provided by the QDNAseq package [72]. Intermediate result files and code is available at Zenodo (https://doi.org/10.5281/zenodo.7945498).

For HMMCopy and PERT, we ran version 0.8.15 of the single-cell pipeline (https://github.com/shahcompbio/single_cell_pipeline) to determine ploidy directly from the aligned bam files with default parameters. We ran the latest version of the Ginkgo platform (https://github.com/robertaboukhalil/ginkgo, version:71da01d9b24b1fcd0deb299b416a0fde676b18f7). For CHISEL[58], we directly use the published results in the case of the 10X Breast tumour dataset. For the ACT samples, we ran CHISEL (v1.1.3) in the *nonormal* mode, with germline SNPs phased using the Michigan Imputation Server. Candidate SNPs were directly called on the merged single-cell bam files. The reference panel used to filter and phase SNPs was TOPMed r2. Phasing was performed using EAGLE2 [80].

For the cell cycle analysis, we ran an early release version of the PERT tool [60] (https://github.com/shahcompbio/scdna_replication_tools, version:7814bdc7572b52a23b7297ff3b67cb89b5e76483).

### Authors’ contributions

MPS: method development, data analysis, manuscript writing; AC, JP, JS, HM, EVO: experimental data collection; AC, AMP, JDB, BC, MG: experimental design and development; GM: supervision, manuscript editing; FM: funding, project conception and oversight, manuscript editing.

## Acknowledgements

We thank Austin Reed, Thomas Bradley and Niko Beerenwinkel as well as all members of the Markowetz and Macintyre labs for helpful discussions and feedback. We thank Simone Zaccaria for help with running CHISEL. We thank the Genomics, Flow Cytometry, Research Instrumentation and Cell Services, and Scientific Computing Core Facilities of the Cancer Research UK Cambridge Institute for technical support. The following plasmids were a gift from Didier Trono: pRSV (Addgene plasmid #12253 ; http://n2t.net/addgene:12253 RRID:Addgene_12253) and pMDLg/pRRE (Addgene plasmid # 12251 ; http://n2t.net/addgene:12251 ; RRID:Addgene_12251)). pVSV-G was a gift from Akitsu Hotta (Addgene plasmid # 138479 ; http://n2t.net/addgene:138479 ; RRID:Addgene_138479). pBOB-EF1-FastFUCCI-Puro was a gift from Kevin Brindle and Duncan Jodrell (Addgene plasmid #86849 ; http://n2t.net/addgene:86849 ; RRID:Addgene_86849)

## Supplementary Materials

**Table S1:**
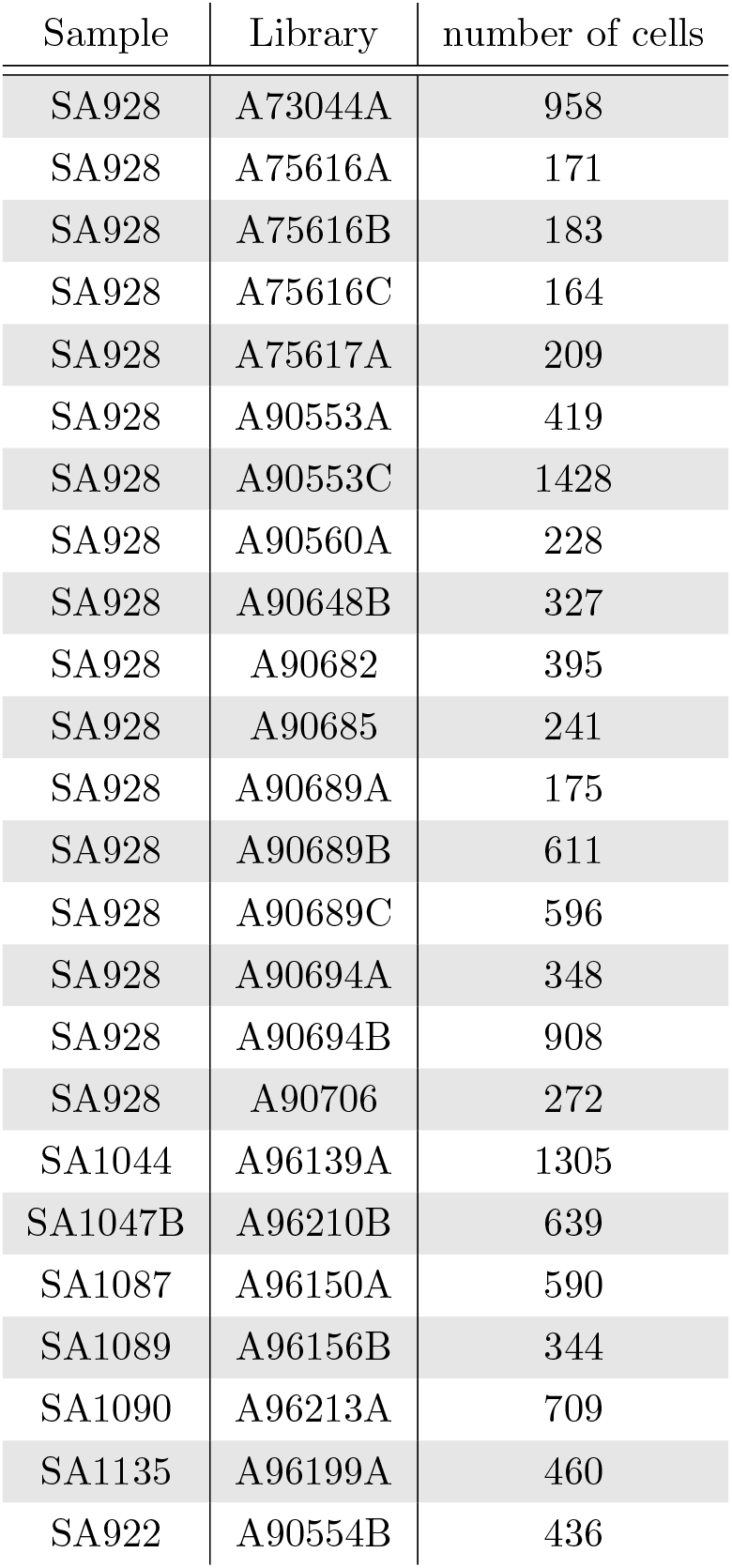
Overview over DLP+ sequencing data used in this study.

| Sample | Library | number of cells |
| --- | --- | --- |
| SA928 | A73044A | 958 |
| SA928 | A75616A | 171 |
| SA928 | A75616B | 183 |
| SA928 | A75616C | 164 |
| SA928 | A75617A | 209 |
| SA928 | A90553A | 419 |
| SA928 | A90553C | 1428 |
| SA928 | A90560A | 228 |
| SA928 | A90648B | 327 |
| SA928 | A90682 | 395 |
| SA928 | A90685 | 241 |
| SA928 | A90689A | 175 |
| SA928 | A90689B | 611 |
| SA928 | A90689C | 596 |
| SA928 | A90694A | 348 |
| SA928 | A90694B | 908 |
| SA928 | A90706 | 272 |
| SA1044 | A96139A | 1305 |
| SA1047B | A96210B | 639 |
| SA1087 | A96150A | 590 |
| SA1089 | A96156B | 344 |
| SA1090 | A96213A | 709 |
| SA1135 | A96199A | 460 |
| SA922 | A90554B | 436 |

**Table S2:** cellenONE isolation parameters for diameter, elongation, circumference, and fluorescence values. Values in parenthesis indicate selection channel (positive/negative).

| Sample | diameter<br>(min,µm) | diameter<br>(max,µm) | Elongation | Transmission | Blue<br>channel | Green<br>channel |
| --- | --- | --- | --- | --- | --- | --- |
| PEO1-FUCCI<br>(G1) | 17 | 24 | 1.8 | 1-255 (+) | 8-255 (+) | NA |
| PEO1-FUCCI<br>(G2) | 20 | 26 | 1.8 | 1-255 (+) | 8-255 (+) | NA |
| PEO1-FUCCI<br>(S) | 19.3 | 23.9 | 1.76 | 1-255 (+) | 8-255 (+) | NA |
| PEO1-PLOIDY | 15.5 | 21 | 1.8 | 1-255 (+) | 8-255 (+) | 5-255 (-) |

**Fig. S1:**
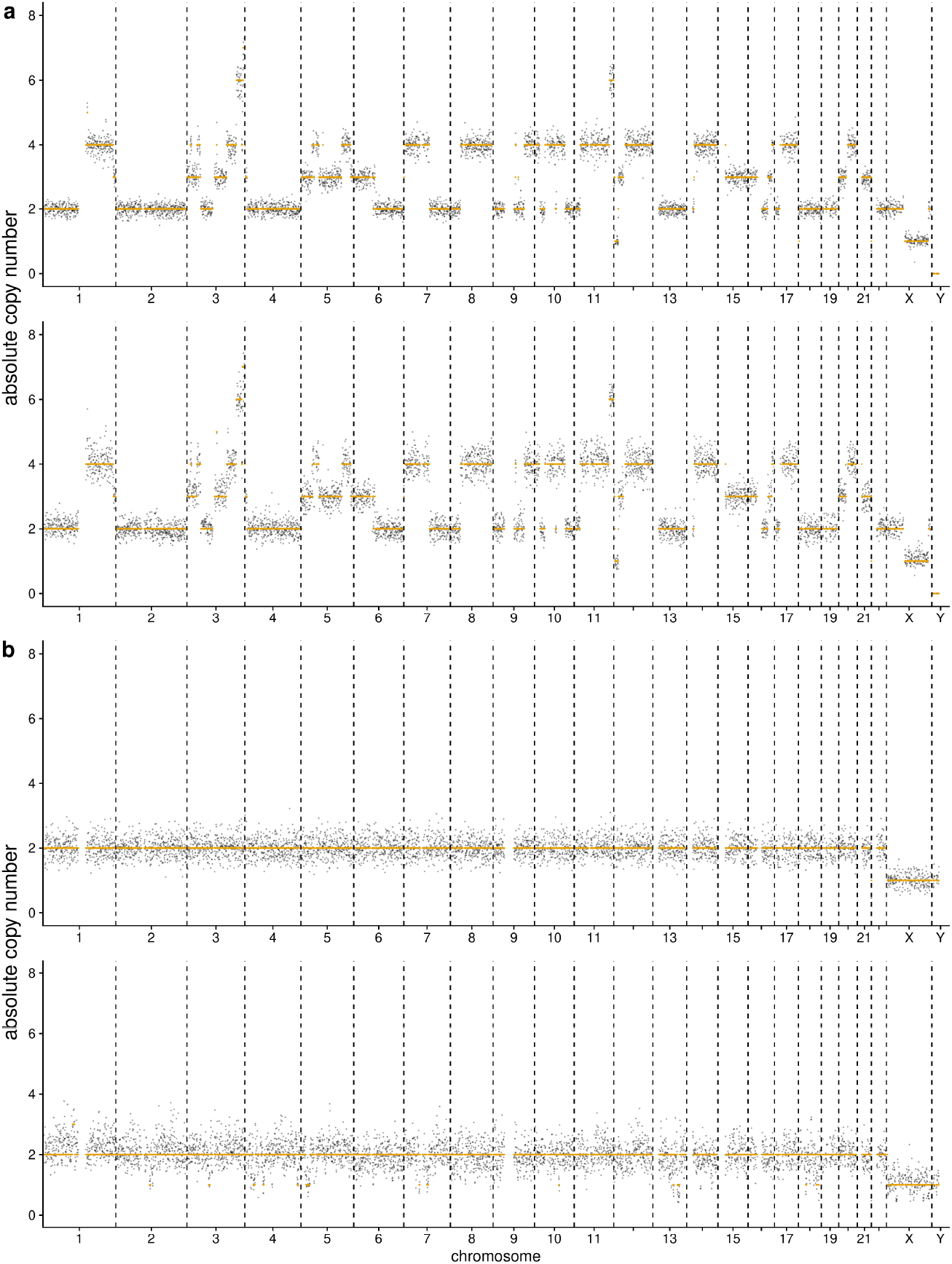
Example cells in G1 and G2 phase of cell cycle. **(a)** Example tumour cell (T-47D) in G1 phase (top panel) and G2 phase (bottom panel) of cell cycle. **(b)** Example normal cell (SA928) in G1 phase (top panel) and G2 phase (bottom panel) of cell cycle. In all cases, cell cycle stage has been verified by DAPI staining based FACS and by subsequent computational analysis. Note, that in case of G2 phase, the initial ploidy solution is off by a multiplicative factor of 2.

**Fig. S2:**
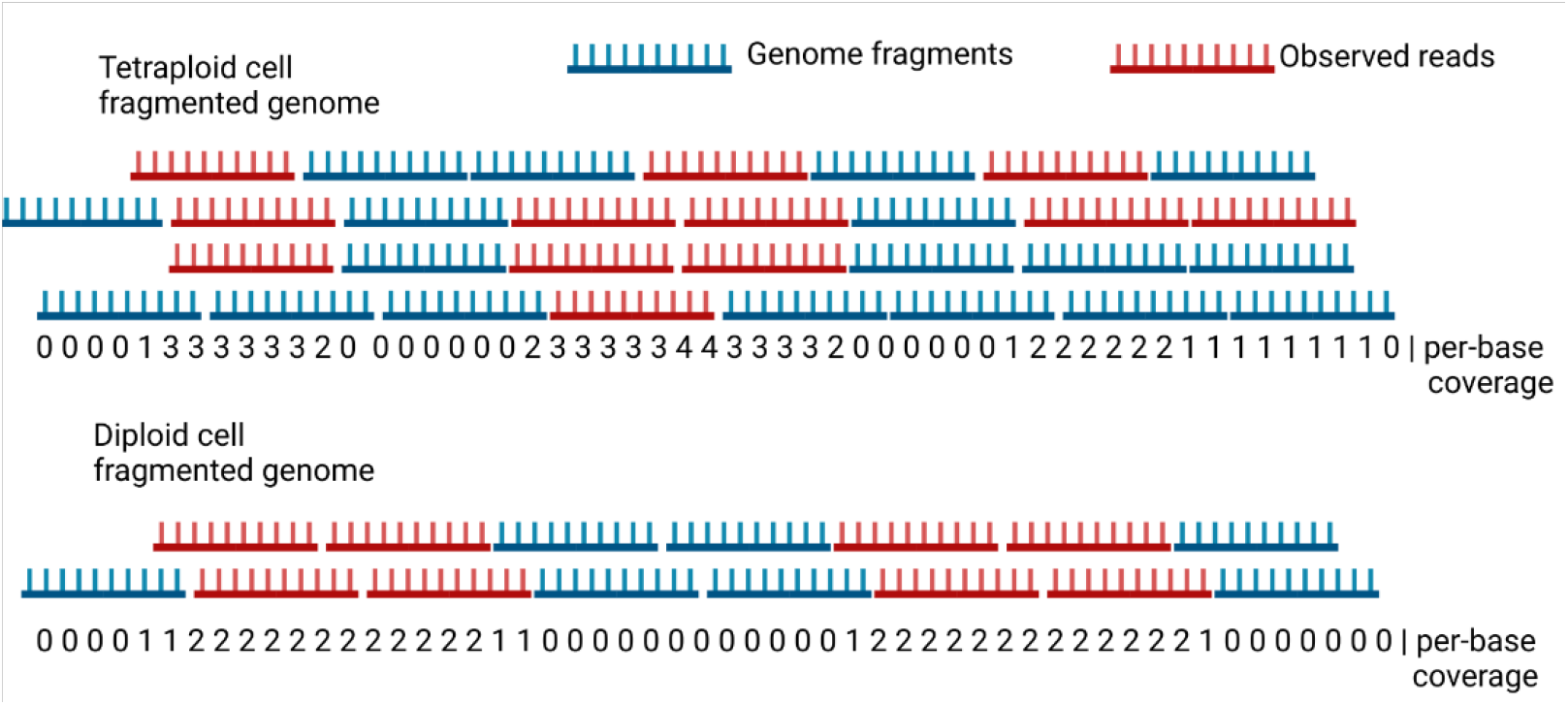
Schematic of read density measure. Read density varies based on the underlying amount of DNA, and the number of overlapping reads can be used to infer the number of original DNA molecules, if the DNA has not been whole-genome amplified. If the DNA is not amplified, each read can be assigned to one strand of DNA in a cell. This makes the ploidy of a cell identifiable by the bound on the number of overlapping reads that can be observed per cell.

**Fig. S3:**
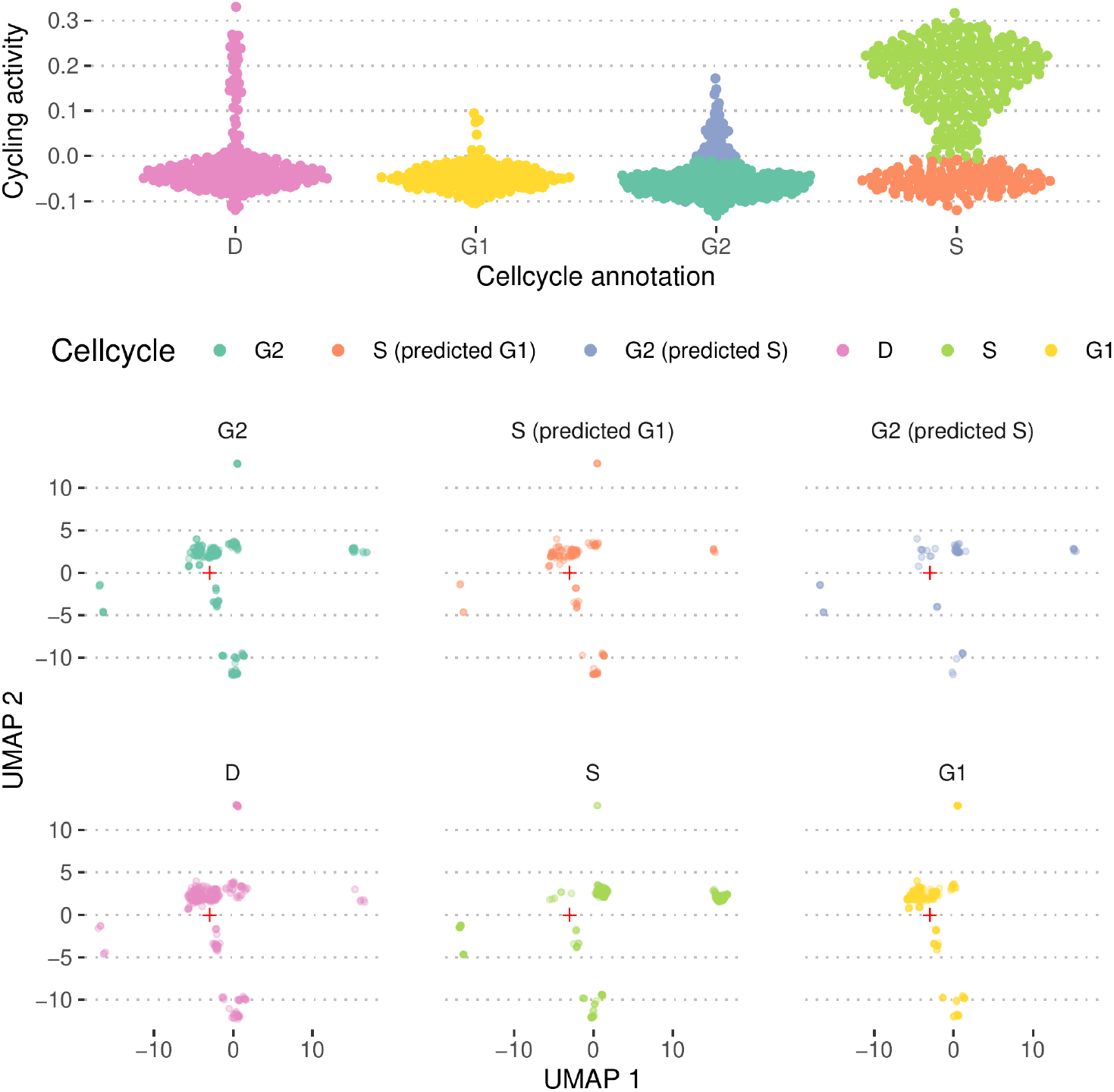
*scAbsolute* predictions for T-47D sample. Top panel shows cycling activity predictions for cells from the T-47D cell line, with DAPI staining based FACS cell cycle annotation on the x-axis. Both for the S phase, and for the G2 phase annotated cells, we observe subgroups that are classified differently by the cycling activity predictor. Looking at copy number profiles in a UMAP representation, we can see that the groups cluster differently based on the cycling activity predictions. Cells that are annotated to be in S phase, but predicted to be in G1 phase (orange), appear to cluster closer with the G1 cells. Similarly, cells in G2 phase that have been predicted to be undergoing replication are clustering more similarly to the cells in S phase. The same pattern can be observed in the raw copy number profiles in Fig. S4.

**Fig. S4:**
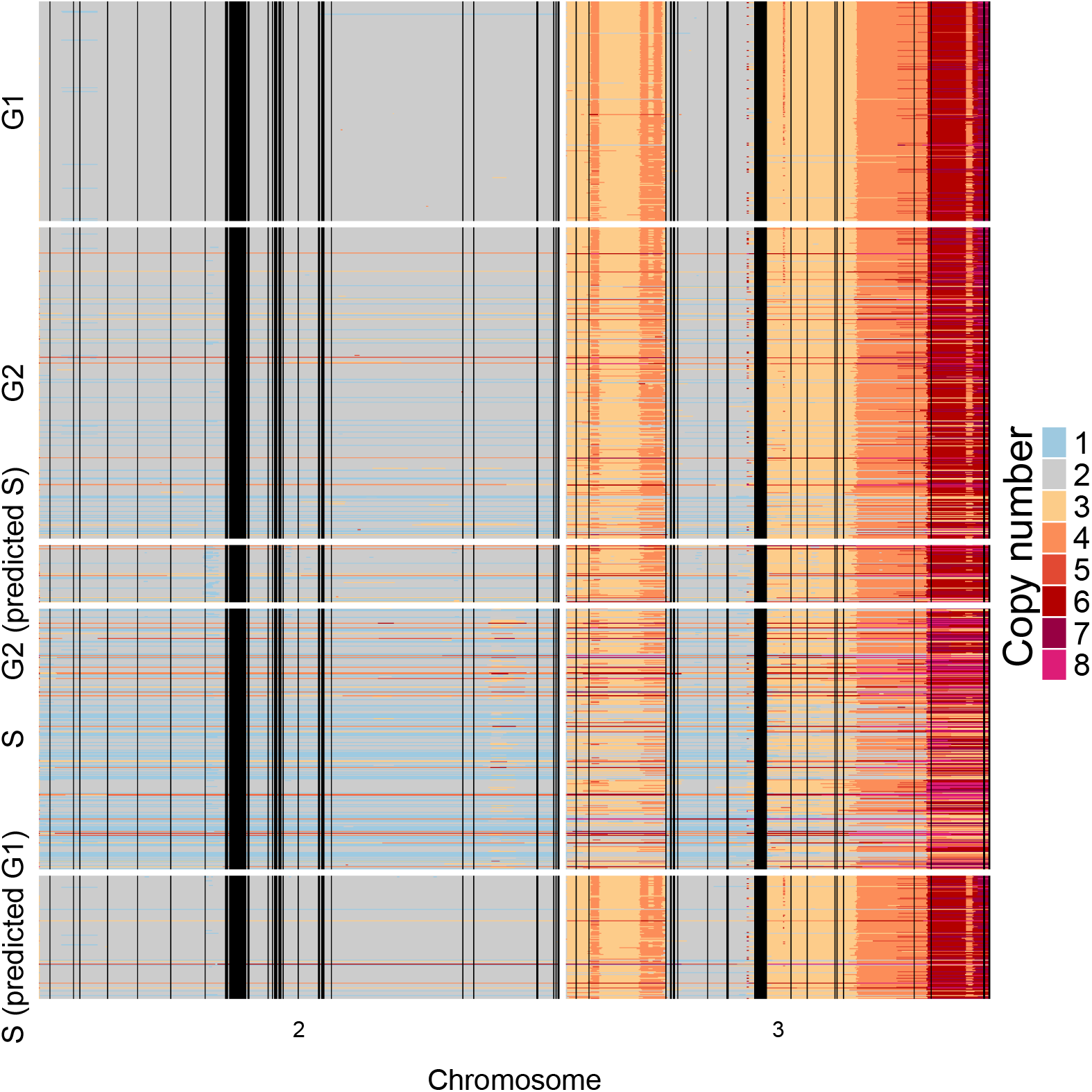
Initial copy number profiles as predicted by scAbsolute for DLP+ T-47D sample for chromosomes 2 and 3. Overall, the vast majority of cells is independently called with the same ploidy. We see that there is a small subset of cells with the wrong ploidy solution and an even smaller group of cells that is otherwise classified wrongly. The algorithm cannot distinguish between G1/G2 cells. We observe some noise, characteristic of S phase cells that are also mentioned in the original publication among the G2 cell population. One can observe a clear difference in copy number profiles between S phase cells predicted to be in G1 phase of the cell cycle, and S phase cells that are predicted to be in S phase based on cycling activity. This might indicate an issue with the underlying ground truth data.

**Fig. S5:**
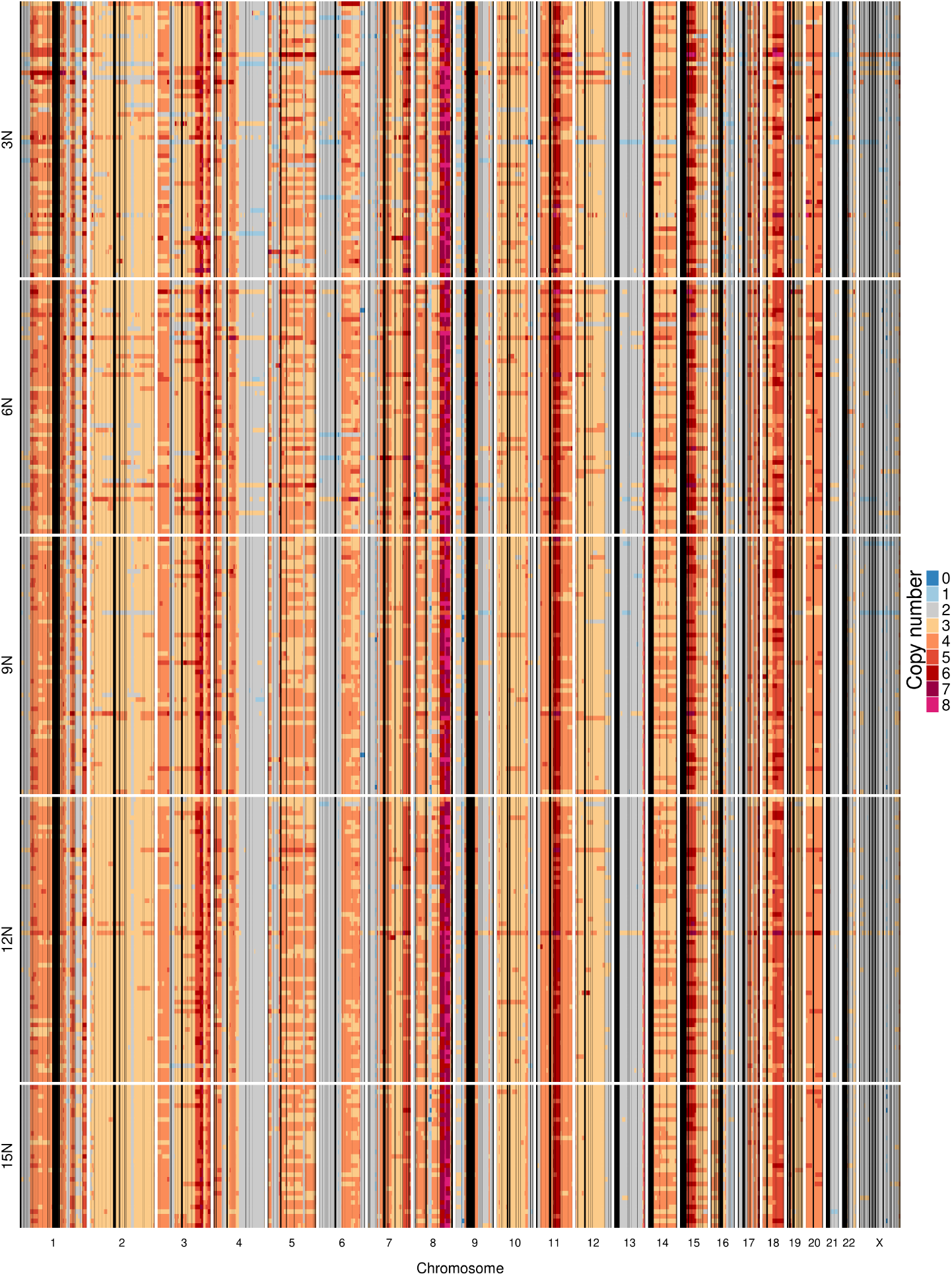
Copy number profiles for PEO1 cell multiplet experiment. Combinations of multiple cells (1-5 cells) are artificially added to the same well and sequenced as a single cell. Here, we enforce a ploidy solution corresponding to the 2N normal state when fitting the copy number profiles. Visual inspection doesn’t allow to distinguish the different ploidy solutions.

**Fig. S6:**
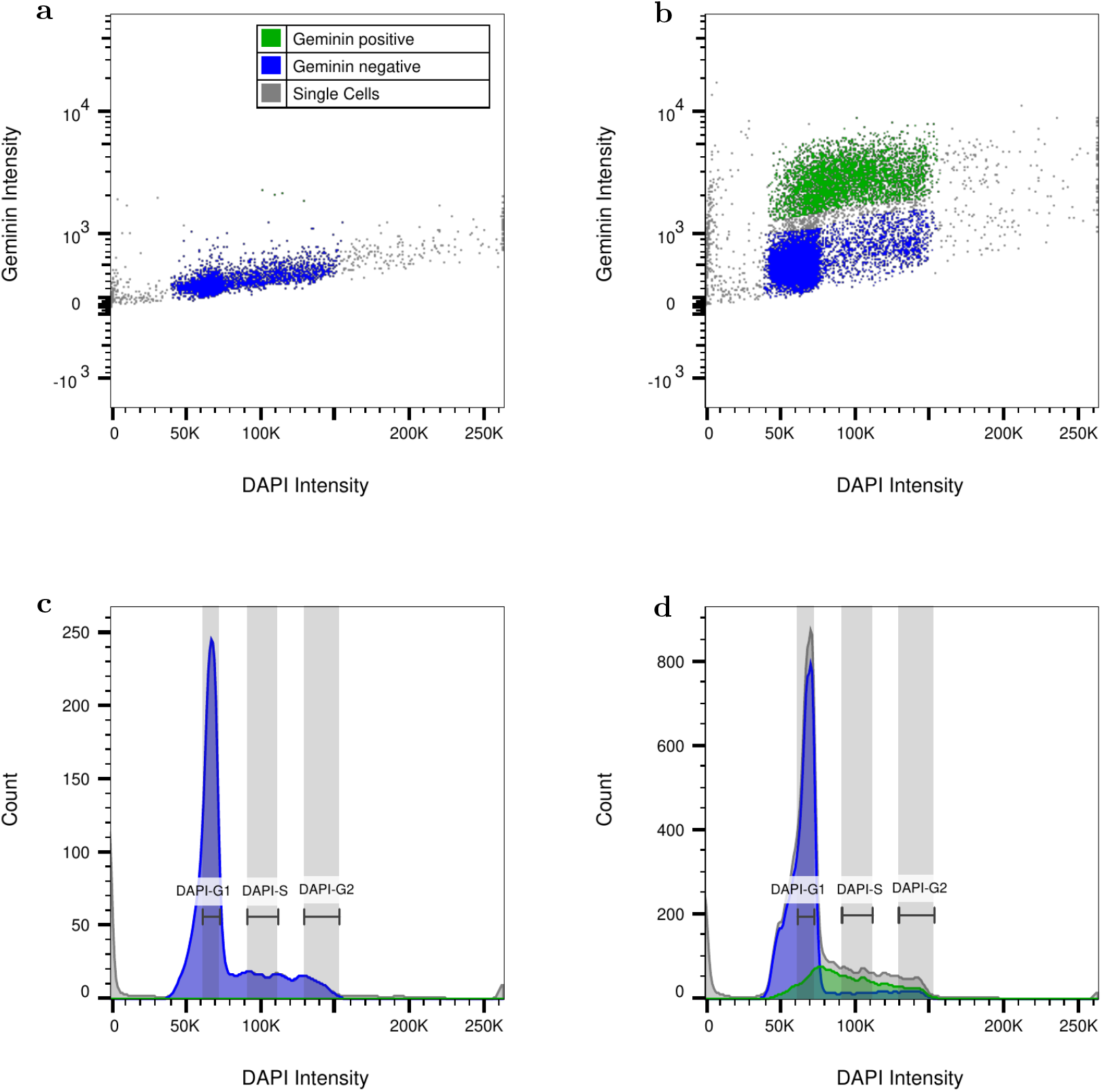
Flow Cytometry Images of normal cells (NA12878) stained with Geminin-AF488 and DAPI for improved G1 cell cycle sorting. Geminin Positive populations – blue; Geminin Negative populations – green. (**a**) Geminin Negative Control - NA12878 stained with DAPI and AF488 secondary antibody. (**b**) NA12878 stained with Geminin/AF488 and DAPI. Manual gating of Geminin positive and Geminin negative populations. (**c**) Cell Cycle curve of Geminin Negative Control using DAPI intensity, with overlay of Geminin gating. DAPI-G1, DAPI-S and DAPI-G2 gating represents original flow cytometry sorting gates using DAPI alone for cell cycle analysis. (**d**) Cell Cycle curve of Geminin stained NA12878 using DAPI intensity. Overlay of Geminin gating reveals Early S phase cells leaking into G1 sorting using DAPI only.

**Fig. S7:**
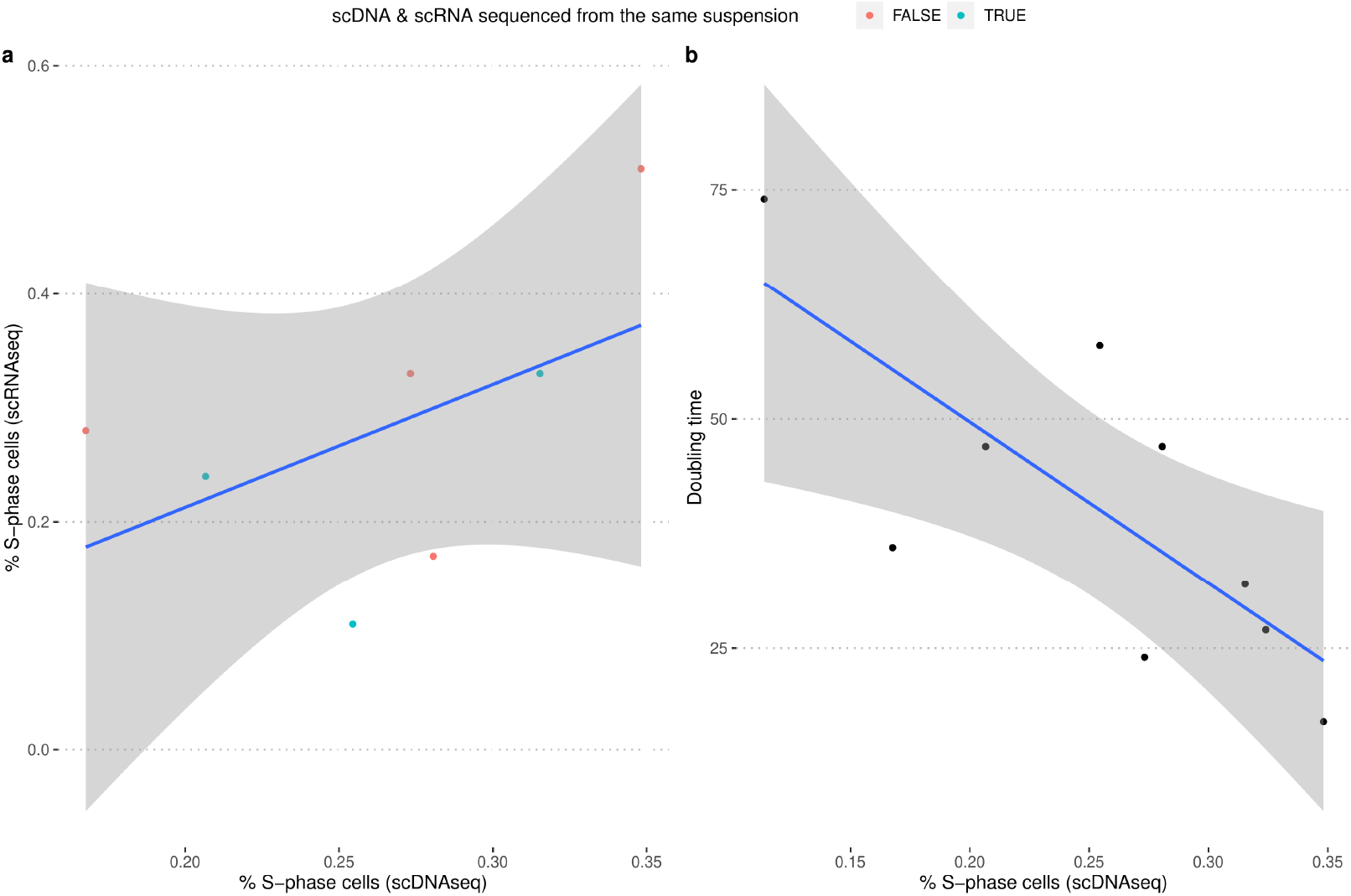
Validation measures for scDNAseq estimates of number of cycling cells in gastric cancer cell lines. **(a)** The number of cycling cells as estimated in scDNAseq data corresponds to estimates of cycling cells based on scRNAseq data. **(b)** Doubling time of cell lines correlates negatively with number of cycling cells as estimated in scDNAseq data.

**Fig. S8:**
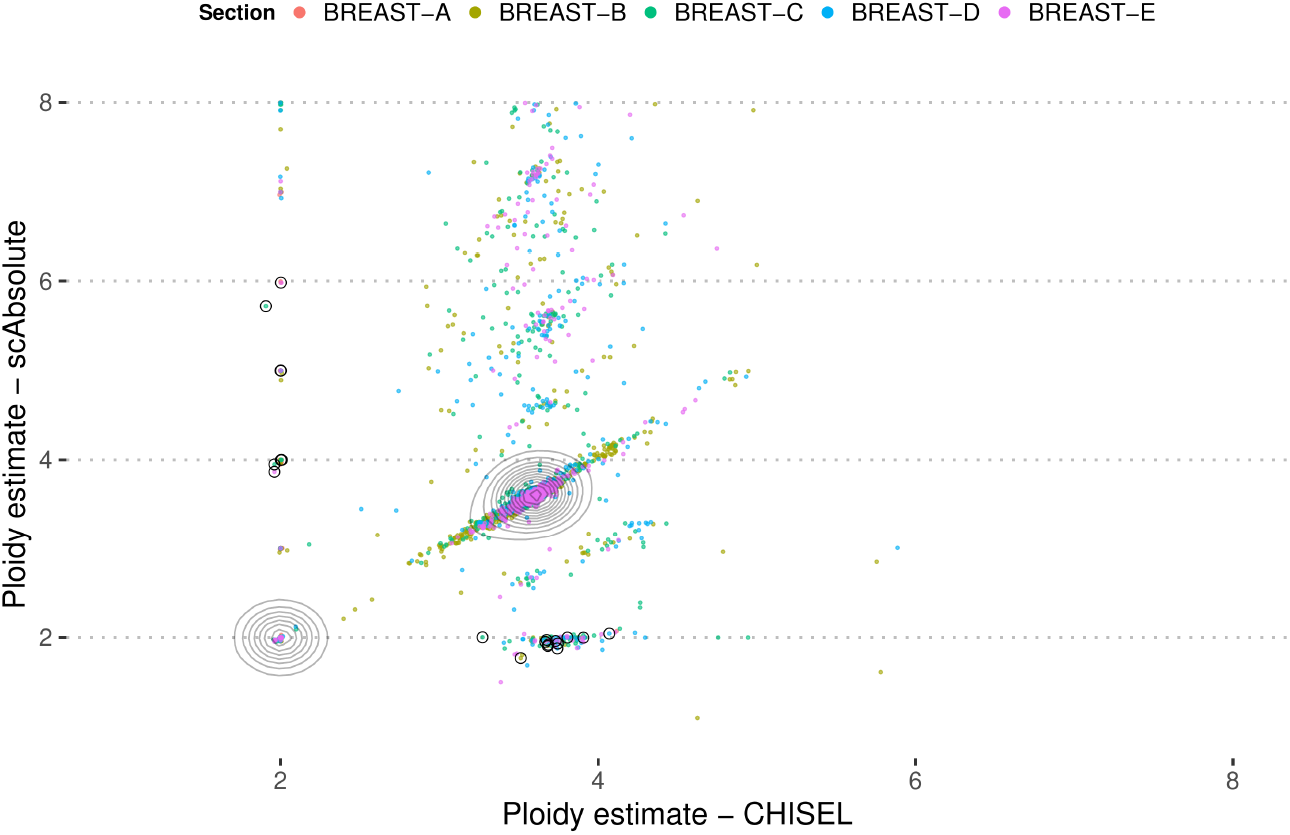
Ploidy predictions for *scAbsolute* and CHISEL on 10X Breast tumour sample. Example cells, shown in Figs. S9 and S10 are marked with black circles. Density areas are indicated on the diagonal, showing a relatively large overlap of predictions in this particular tumour sample.

**Fig. S9:**
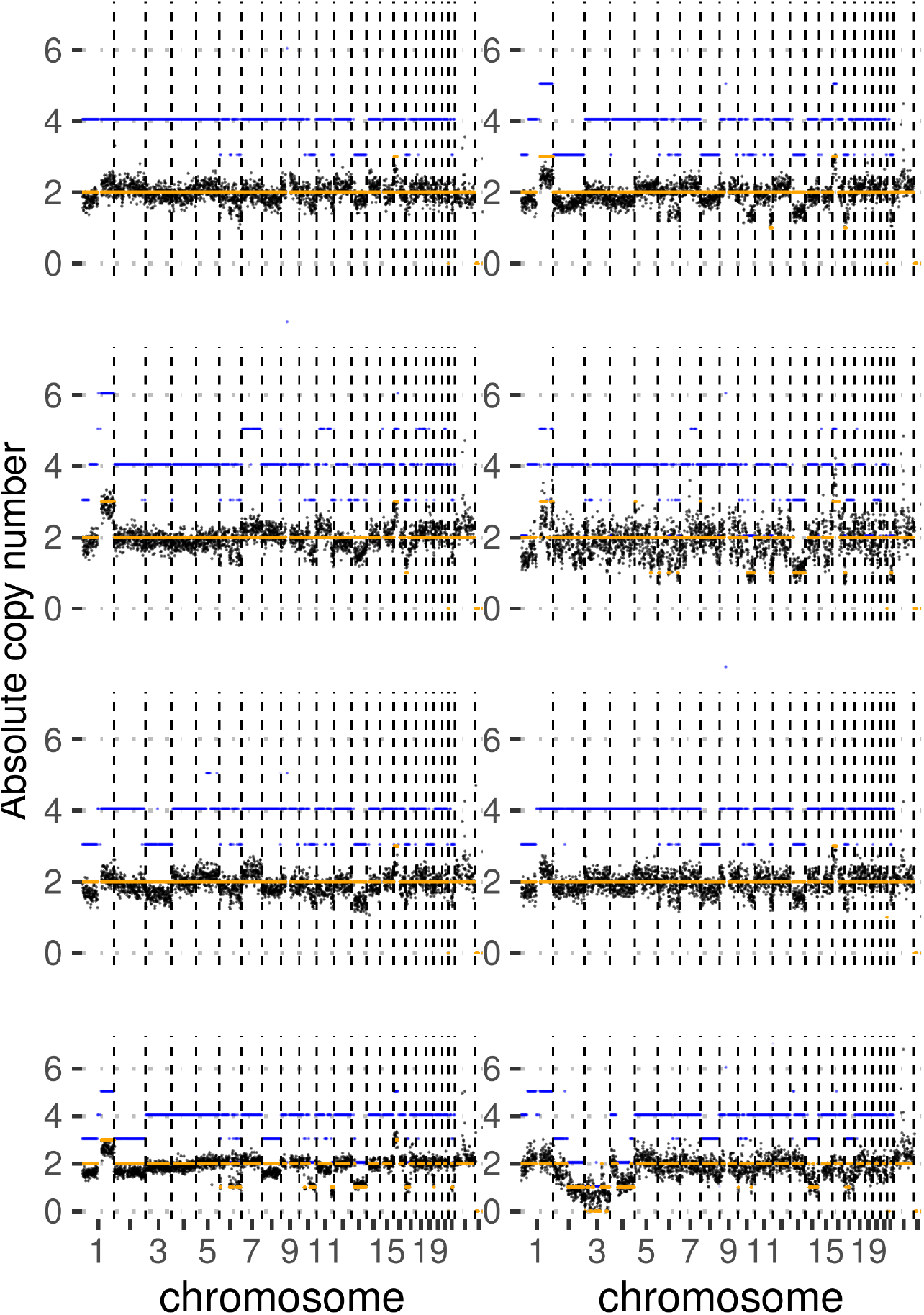
Randomly selected copy number profiles, for which *scAbsolute* predicts a diploid copy number profile and CHISEL has a contradictory prediction. *scAbsolute* predictions are shown in orange, and CHISEL in blue. In some cases it appears that *scAbsolute* fails to detect slight copy number changes, indicating a more conservative segmentation. CHISEL tends to select higher ploidy solutions that might not necessarily be the minimum ploidy fit in these cases, as shown in the bottom row.

**Fig. S10:**
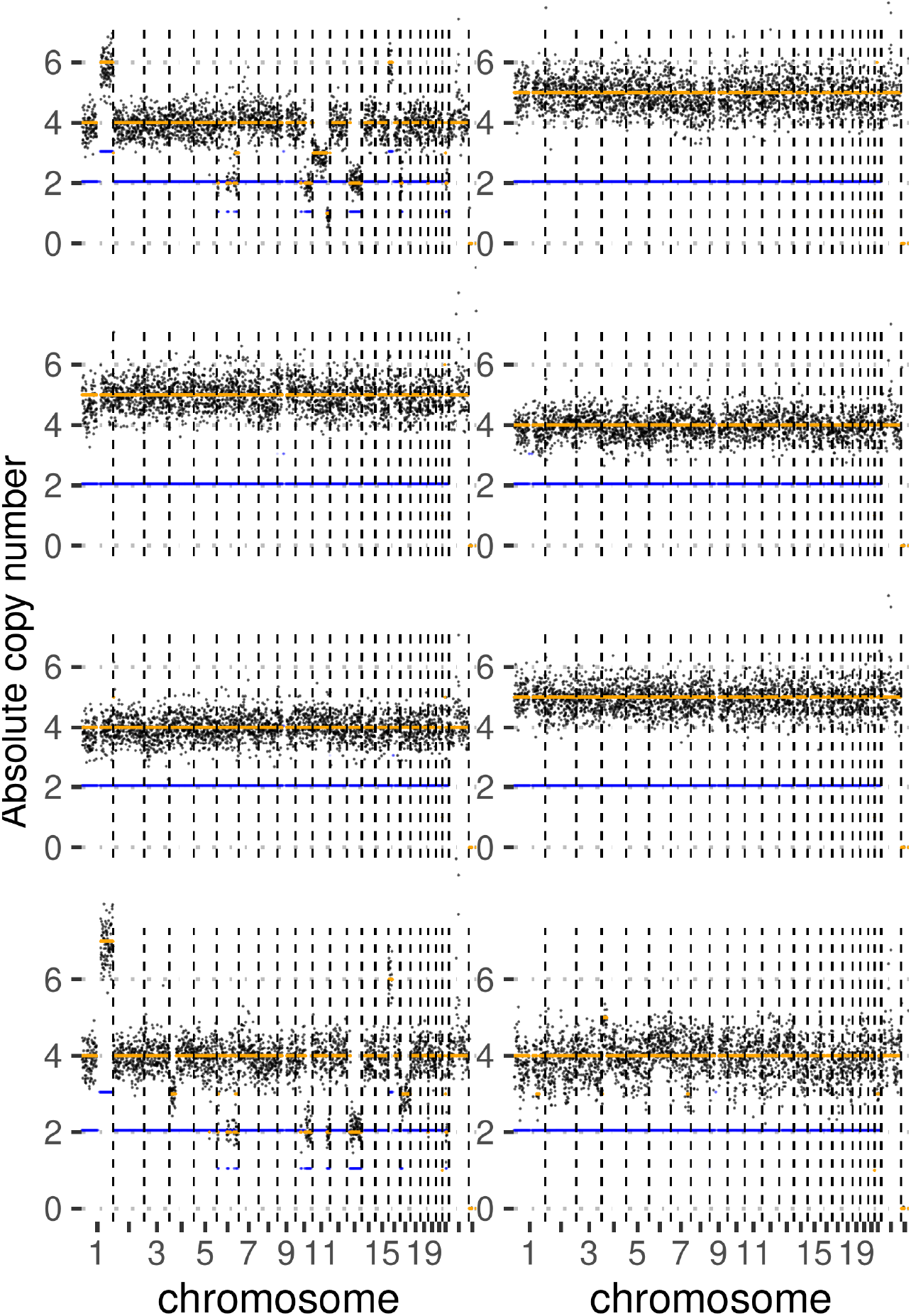
Randomly selected copy number profiles, for which CHISEL predicts a diploid copy number profile and *scAbsolute* has a contradictory prediction. *scAbsolute* predictions are shown in orange, and CHISEL in blue. Here, *scAbsolute* selects potentially wrong solutions in cases with very minor erroneous copy number changes. Note that the number of these cases is very small (about 1% of cells). At the same time, it appears more accurate in cases with multiple levels of copy number changes (top left and bottom left examples).

**Fig. S11:**
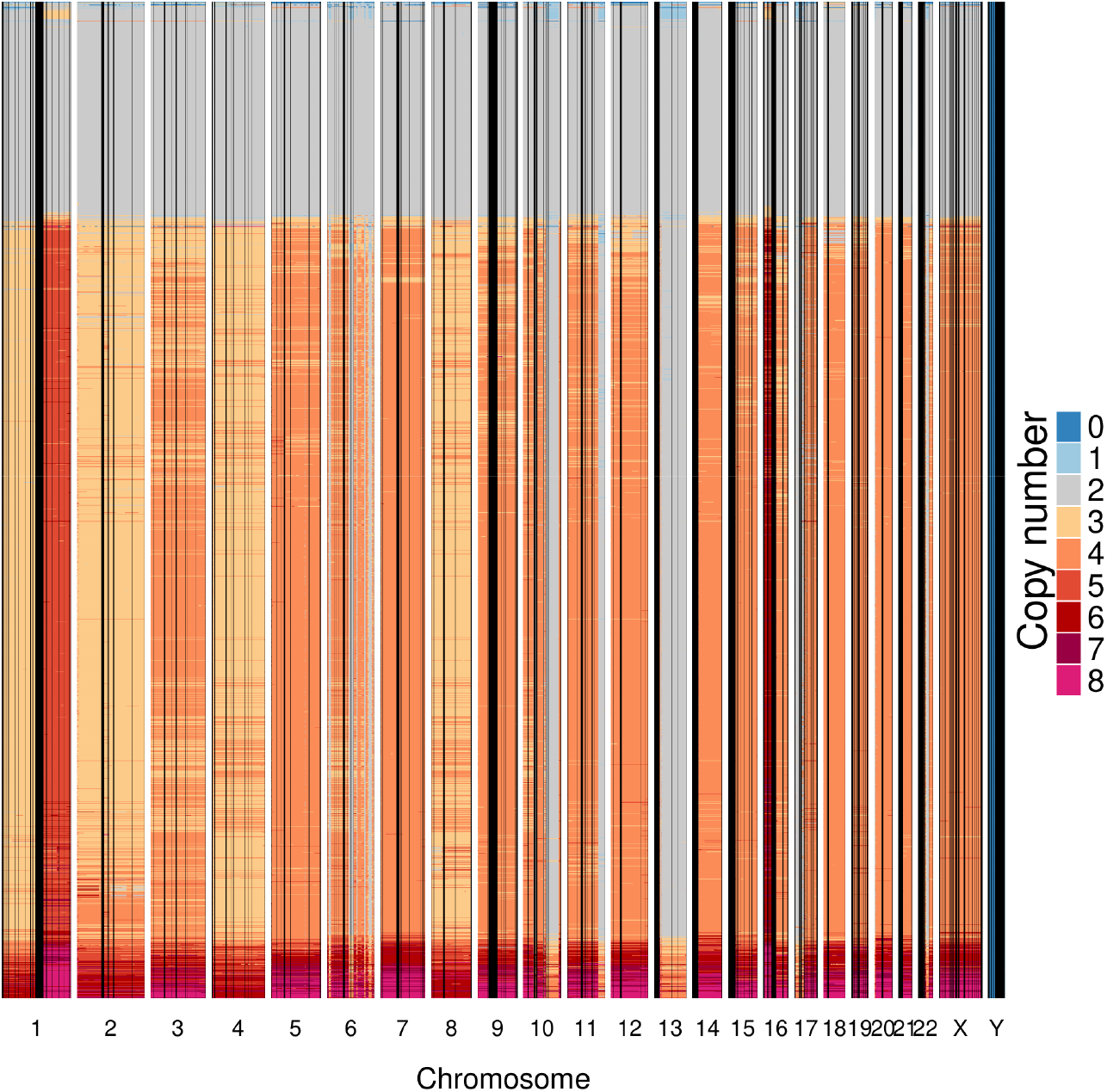
Copy number prediction for Patient S0 (section E) using *scAbsolute*. The prediction reflects the general copy number landscape for the sample as presented in Zaccaria and Raphael [58]. Note that the cells have not been quality controlled, and this explains the small number of ploidy outliers at high and low ploidies. Overall it appears relatively easy to detect higher ploidy states in this dataset, given the number of copy number segments at varying ploidy levels. This might be indicative of a relatively early WGD event leading to the observed copy number profiles.

**Fig. S12:**
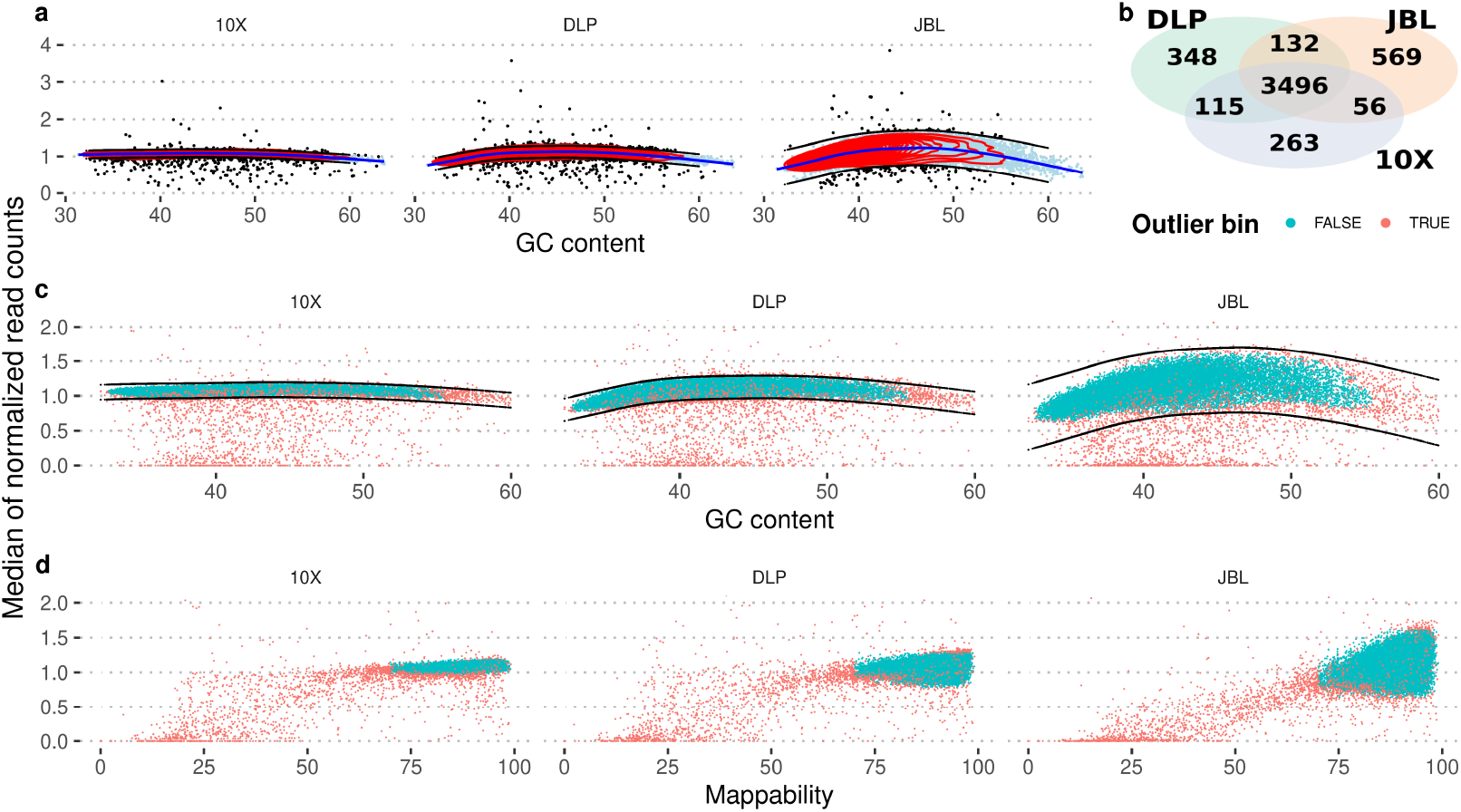
Bin level quality control for autosomes across sequencing technologies. **a)** Generative additive model smoothing (blue line) and kernel density estimation (red contours) to identify genomic bins that have below or above average median expected read counts. **b)** Number of genomic bins identified as outliers by sequencing technology. We remove the union of all outliers. **c+d)** Genomic bins identified as outliers (in red) across different sequencing technologies as a function of GC content **(c)**, and mappability **(d)**.

**Fig. S13:**
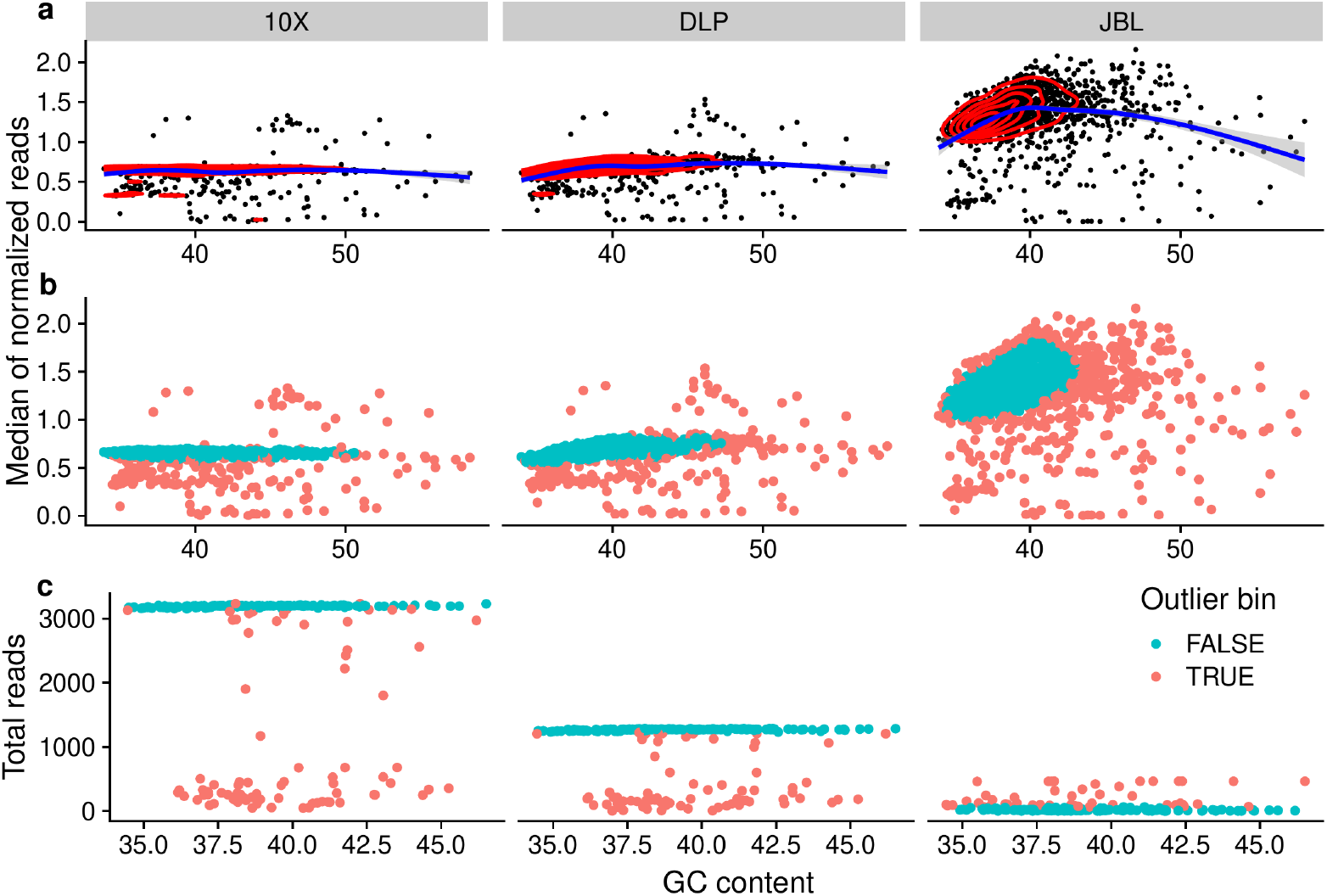
Bin level quality control for sex chromosomes across sequencing technologies. **a)** Generative additive model smoothing (blue line) and kernel density estimation (red contours) to identify genomic bins that have below or above average median expected read counts for the X chromosome. **b)** Outlier bins that are removed based on the fits in a) are marked. **c)** Total (absolute) reads per bin observed on the Y chromosome. Outlier bins (based on deviation from median number of total reads) are marked in red.

## Supplementary methods

### 0.1 Derivation of Dirichlet Process Gaussian Mixture Model for scAbsolute algorithm

We implement a DPGMM model, following [67]. We assume throughout the document, that the dimensionality of our problem is 1. We use the mean-field approximation to estimate 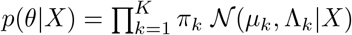. The model is depicted below. Note that we restrict the means *μ*_*k*_ = *ξ · k. K* refers to the number of components obtained after fitting the model (*T* is the maximum possible number), and *π*_*k*_ refers to the normalized weight of the components.

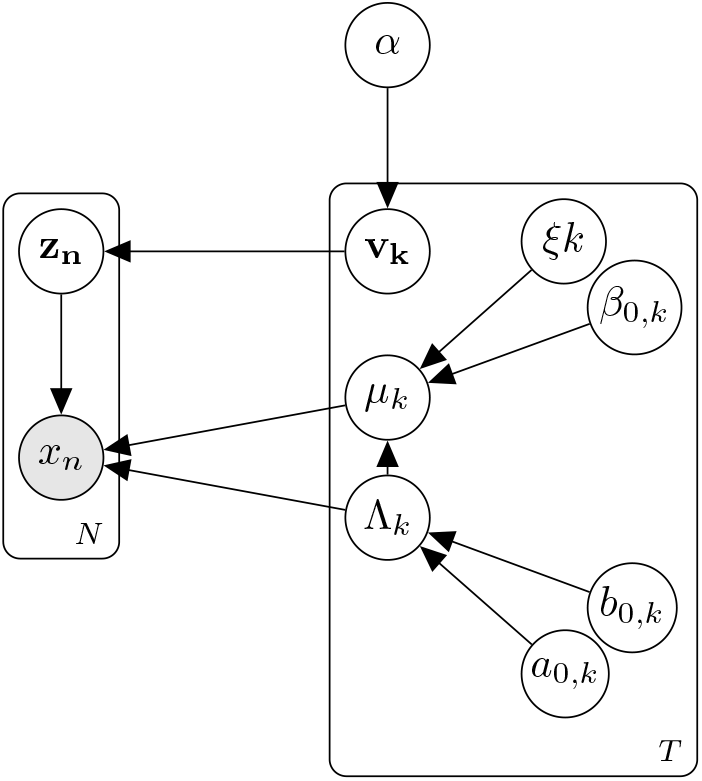

#### 0.1.1 Model and priors

Mean-field approximation

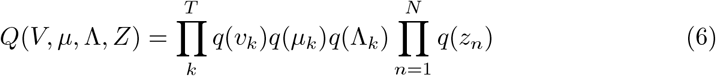

Prior distributions.

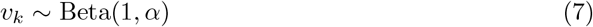

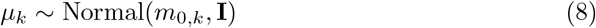

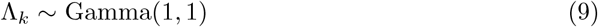

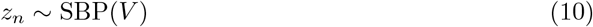

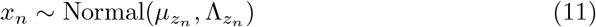

Variational distributions.

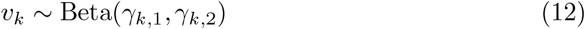

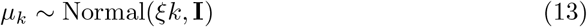

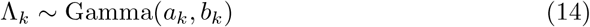

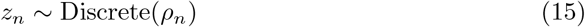

#### 0.1.2 Variational bound

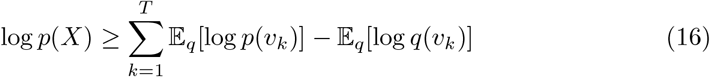

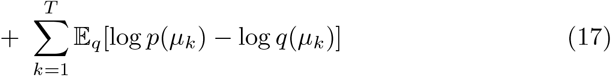

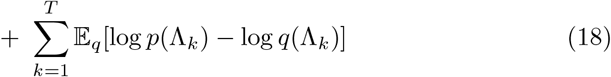

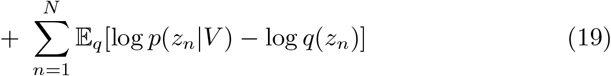

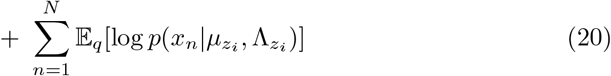

In the following, we derive the terms in the variational bound.

*v*_*k*_ ***terms***

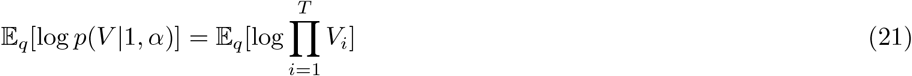

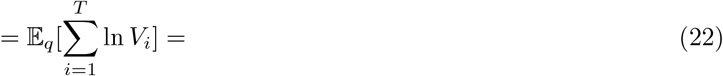

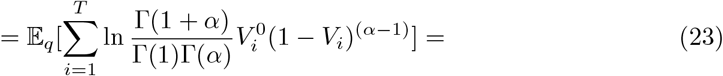

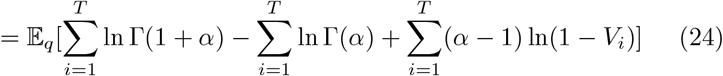

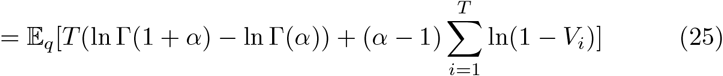

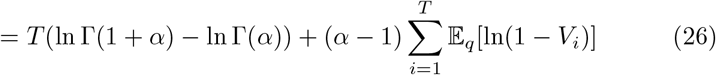

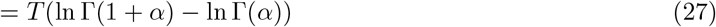

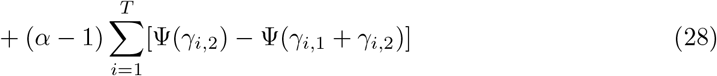

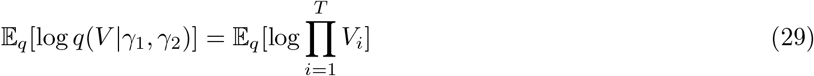

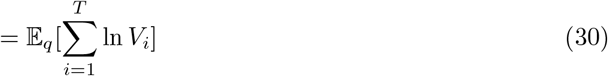

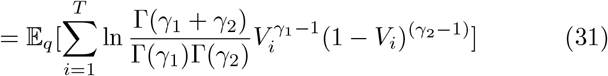

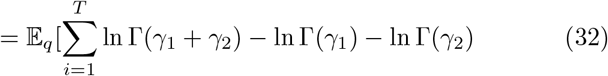

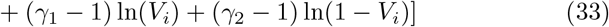

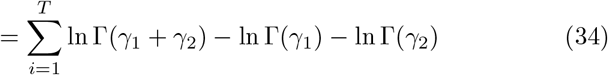

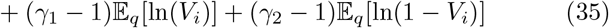

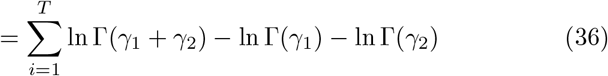

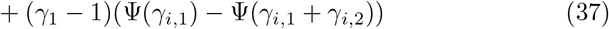

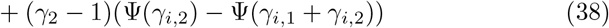

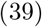

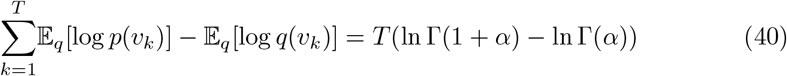

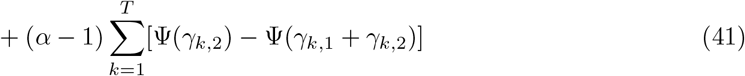

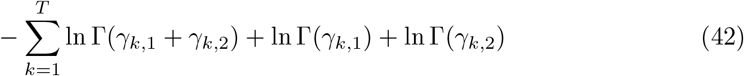

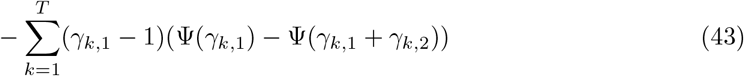

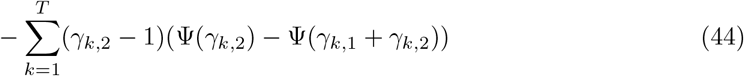

*μ*_*k*_ ***terms***

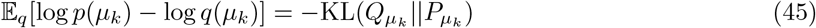

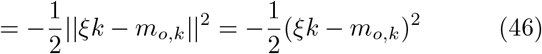

Λ_*k*_ ***terms***

We use the inverse scale parameter characterization of the Gamma distribution, with Λ_*k*_ ∼ *𝔾*(Λ_*k*_|*a*_*k*,0_, *b*_*k*,0_), for *a*_*k*,0_, *b*_*k*,0_ = 1, and 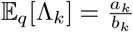.

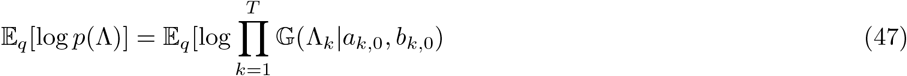

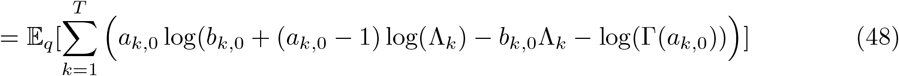

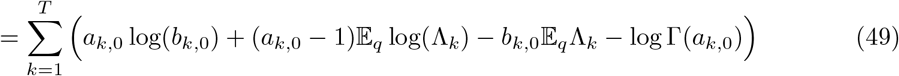

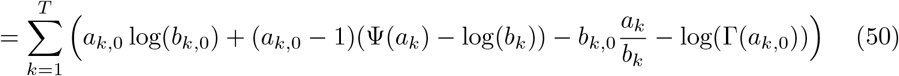

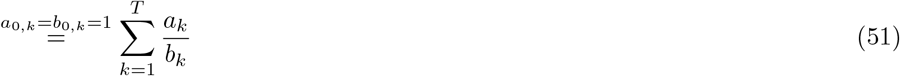

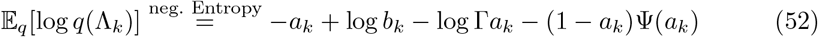

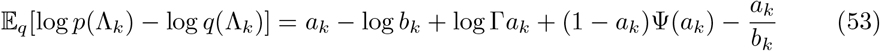

*z*_*n*_ ***terms***

Here, we use the equations as presented by [67, p. 129].

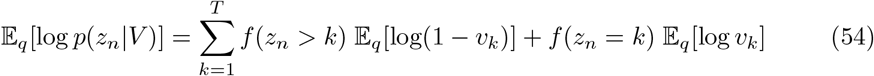

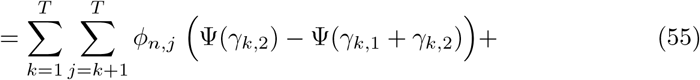

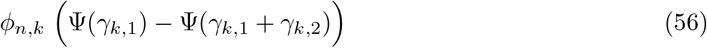

with the following definitions

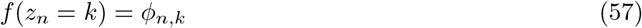

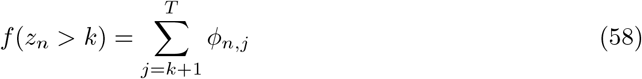

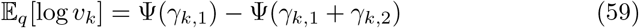

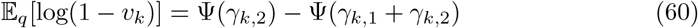

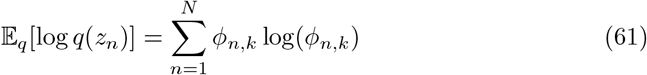

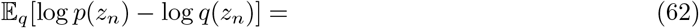

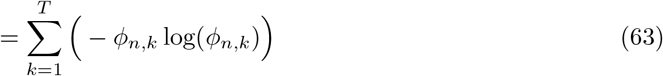

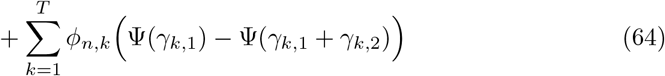

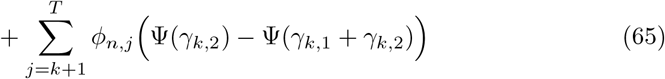

Derivation: (note this leads to a slightly different result than presented in Blei.)

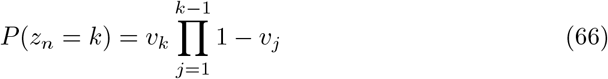

We use the following identities of the Beta distribution: for *B* ∼ Beta(*b*_1_, *b*_2_)

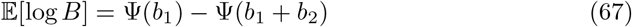

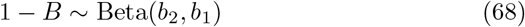

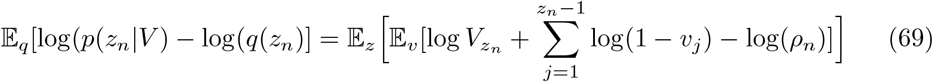

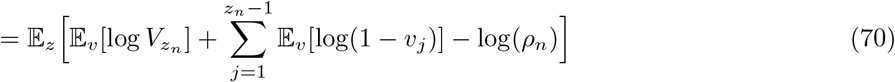

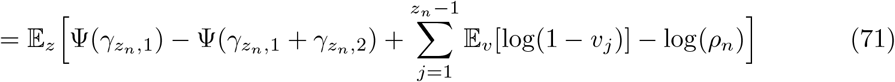

*x*_*n*_ ***terms***

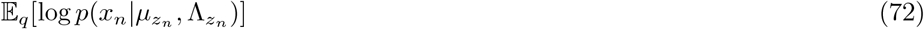

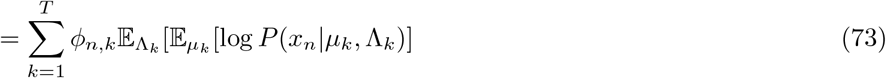

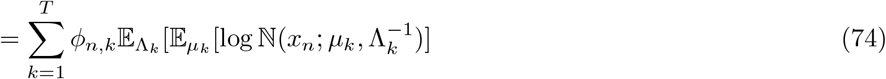

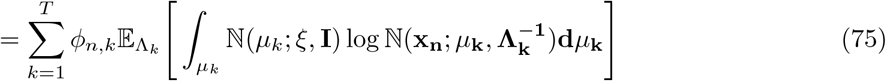

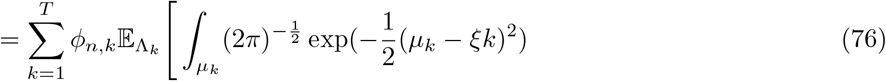

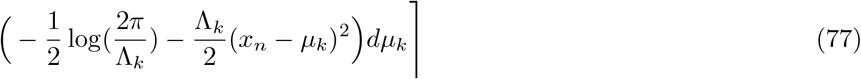

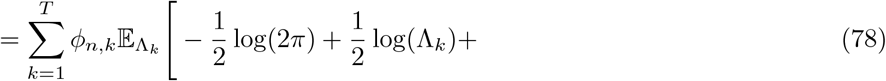

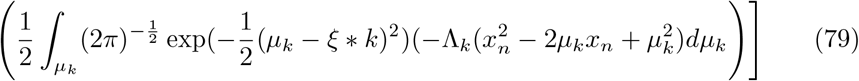

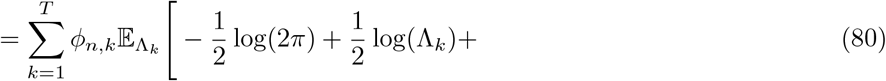

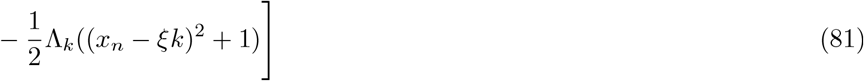

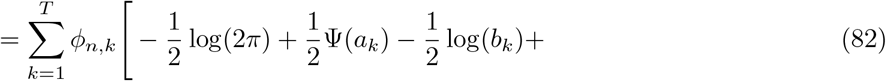

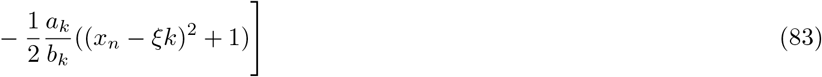

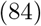

We will refer to the following term later on

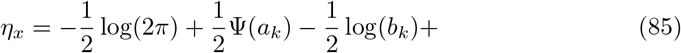

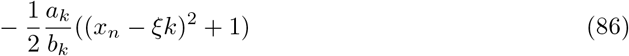

#### 0.1.3 Variational updates

##### Updates for v_k_

These update equations are given in [67, p. 129].

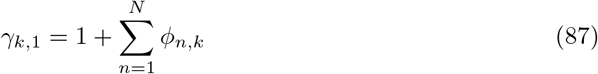

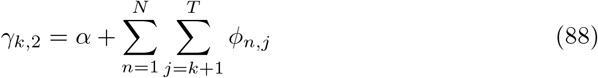

##### Updates for ξ

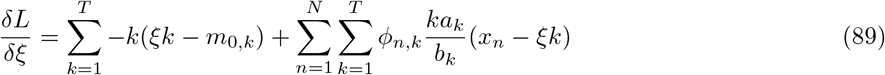

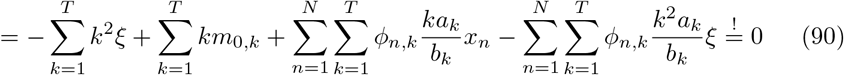

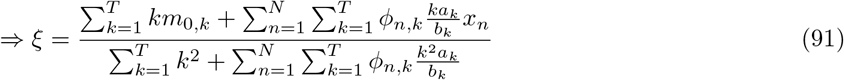

##### Updates for a_k_ and b_k_

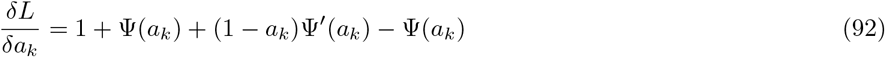

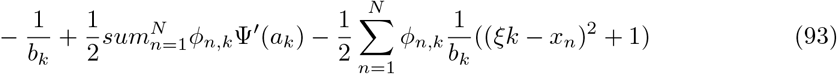

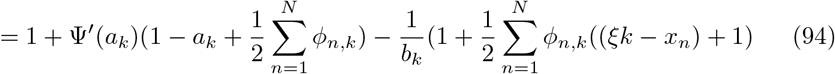

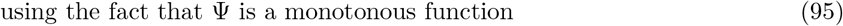

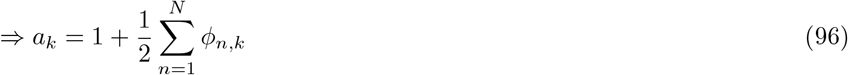

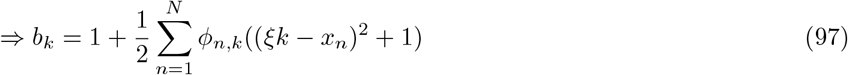

Computing the partial derivative 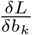 and setting the values of *a*_*k*_ and *b*_*k*_ to the above results satisfies the condition.

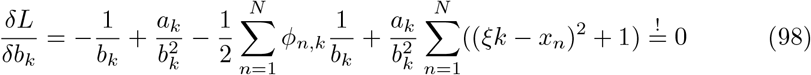

##### Updates for ρ_n_

Here, we need to take into account the constraint 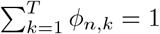 We do this by using Lagrange multipliers.

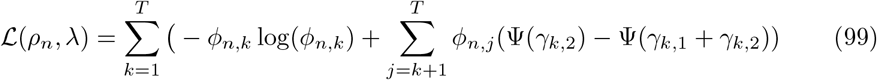

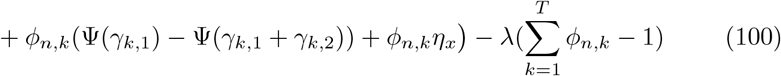

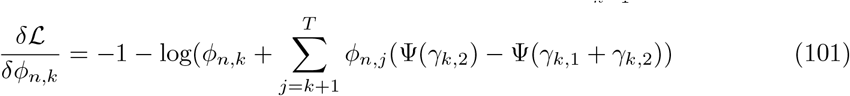

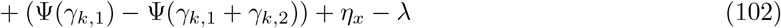

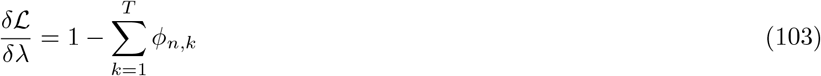

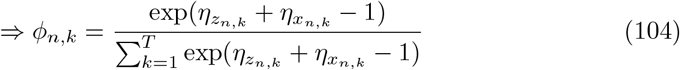

## References

[1] Zack TI, Schumacher SE, Carter SL, Cherniack AD, Saksena G, Tabak B, et al. Pan-Cancer Patterns of Somatic Copy Number Alteration. Nature Genetics. 2013;45(10):1134–1140. https://doi.org/10.1038/ng.2760.

[2] Lukow DA, Sheltzer JM. Chromosomal Instability and Aneuploidy as Causes of Cancer Drug Resistance. Trends in Cancer. 2021;https://doi.org/10.1016/j.trecan.2021.09.002.

[3] McGranahan N, Swanton C. Biological and Therapeutic Impact of Intratumor Heterogeneity in Cancer Evolution. Cancer Cell. 2015;27(1):15–26. https://doi.org/10.1016/j.ccell.2014.12.001.

[4] Beroukhim R, Mermel CH, Porter D, Wei G, Raychaudhuri S, Donovan J, et al. The Landscape of Somatic Copy-Number Alteration across Human Cancers. Nature. 2010;463(7283):899–905. https://doi.org/10.1038/nature08822.

[5] Ciriello G, Miller ML, Aksoy BA, Senbabaoglu Y, Schultz N, Sander C. Emerging Landscape of Oncogenic Signatures across Human Cancers. Nature Genetics. 2013;45(10):1127–1133. https://doi.org/10.1038/ng.2762.

[6] Taylor AM, Shih J, Ha G, Gao GF, Zhang X, Berger AC, et al. Genomic and Functional Approaches to Understanding Cancer Aneuploidy. Cancer Cell. 2018;33(4):676–689.e3. https://doi.org/10.1016/j.ccell.2018.03.007.

[7] Chowdhury SA, Shackney SE, Heselmeyer-Haddad K, Ried T, Schäffer AA, Schwartz R. Algorithms to Model Single Gene, Single Chromosome, and Whole Genome Copy Number Changes Jointly in Tumor Phylogenetics. PLOS Computational Biology. 2014;10(7):e1003740. https://doi.org/10.1371/journal.pcbi.1003740.

[8] Schwarz RF, Trinh A, Sipos B, Brenton JD, Goldman N, Markowetz F. Phlogenetic Quantification of Intra-tumour Heterogeneity. PLOS Computational Biology. 2014;10(4):e1003535. https://doi.org/10.1371/journal.pcbi.1003535.

[9] Burrell RA, McGranahan N, Bartek J, Swanton C. The Causes and Consequences of Genetic Heterogeneity in Cancer Evolution. Nature. 2013;501(7467):338–345. https://doi.org/10.1038/nature12625.

[10] Sansregret L, Vanhaesebroeck B, Swanton C. Determinants and Clinical Implications of Chromosomal Instability in Cancer. Nature Reviews Clinical Oncology. 2018;15(3):139–150. https://doi.org/10.1038/nrclinonc.2017.198.

[11] Bakhoum SF, Danilova OV, Kaur P, Levy NB, Compton DA. Chromosomal Instability Substantiates Poor Prognosis in Patients with Diffuse Large B-cell Lymphoma. Clinical Cancer Research. 2011;17(24):7704–7711. https://doi.org/10.1158/1078-0432.CCR-11-2049.

[12] Davoli T, Uno H, Wooten EC, Elledge SJ. Tumor Aneuploidy Correlates with Markers of Immune Evasion and with Reduced Response to Immunotherapy. Science. 2017;355(6322):eaaf8399. https://doi.org/10.1126/science.aaf8399.

[13] Buccitelli C, Salgueiro L, Rowald K, Sotillo R, Mardin BR, Korbel JO. Pan-Cancer Analysis Distinguishes Transcriptional Changes of Aneuploidy from Proliferation. Genome Research. 2017;27(4):501–511. https://doi.org/10.1101/gr.212225.116.

[14] López S, Lim EL, Horswell S, Haase K, Huebner A, Dietzen M, et al. Interplay between Whole-Genome Doubling and the Accumulation of Deleterious Alterations in Cancer Evolution. Nature Genetics. 2020;52(3):283–293. https://doi.org/10.1038/s41588-020-0584-7.

[15] Goupil A, Nano M, Letort G, Gemble S, Edwards F, Goundiam O, et al. Chromosomes Function as a Barrier to Mitotic Spindle Bipolarity in Polyploid Cells. Journal of Cell Biology. 2020;219(4):e201908006. https://doi.org/10.1083/jcb.201908006.

[16] Gemble S, Wardenaar R, Keuper K, Srivastava N, Nano M, Macé AS, et al. Genetic Instability from a Single S Phase after Whole-Genome Duplication. Nature. 2022;604(7904):146–151. https://doi.org/10.1038/s41586-022-04578-4.

[17] Storchova Z, Pellman D. From Polyploidy to Aneuploidy, Genome Instability and Cancer. Nature Reviews Molecular Cell Biology. 2004;5(1):45–54. https://doi.org/10.1038/nrm1276.

[18] Storchova Z, Kuffer C. The Consequences of Tetraploidy and Aneuploidy. Journal of Cell Science. 2008;121(23):3859–3866. https://doi.org/10.1242/jcs.039537.

[19] Fujiwara T, Bandi M, Nitta M, Ivanova EV, Bronson RT, Pellman D. Cytokinesis Failure Generating Tetraploids Promotes Tumorigenesis in P53-Null Cells. Nature. 2005;437(7061):1043–1047. https://doi.org/10.1038/nature04217.

[20] Dewhurst SM, McGranahan N, Burrell RA, Rowan AJ, Grönroos E, Endesfelder D, et al. Tolerance of Whole-Genome Doubling Propagates Chromosomal Instability and Accelerates Cancer Genome Evolution. Cancer Discovery. 2014;4(2):175–185. https://doi.org/10.1158/2159-8290.CD-13-0285.

[21] Ganem NJ, Godinho SA, Pellman D. A Mechanism Linking Extra Centrosomes to Chromosomal Instability. Nature. 2009;460(7252):278–282. https://doi.org/10.1038/nature08136.

[22] Quinton RJ, DiDomizio A, Vittoria MA, Kotynkova K, Ticas CJ, Patel S, et al. Whole-Genome Doubling Confers Unique Genetic Vulnerabilities on Tumour Cells. Nature. 2021;590(7846):492–497. https://doi.org/10.1038/s41586-020-03133-3.

[23] Carter SL, Cibulskis K, Helman E, McKenna A, Shen H, Zack T, et al. Absolute Quantification of Somatic DNA Alterations in Human Cancer. Nature Biotechnology. 2012;30(5):413–421. https://doi.org/10.1038/nbt.2203.

[24] Dentro SC, Leshchiner I, Haase K, Tarabichi M, Wintersinger J, Deshwar AG, et al. Characterizing Genetic Intra-Tumor Heterogeneity across 2,658 Human Cancer Genomes. Cell. 2021;184(8):2239–2254.e39. https://doi.org/10.1016/j.cell.2021.03.009.

[25] Bielski CM, Zehir A, Penson AV, Donoghue MTA, Chatila W, Armenia J, et al. Genome Doubling Shapes the Evolution and Prognosis of Advanced Cancers. Nature Genetics. 2018;50(8):1189–1195. https://doi.org/10.1038/s41588-018-0165-1.

[26] Baslan T, Morris JP, Zhao Z, Reyes J, Ho YJ, Tsanov KM, et al. Ordered and Deterministic Cancer Genome Evolution after P53 Loss. Nature. 2022;608(7924):795–802. https://doi.org/10.1038/s41586-022-05082-5.

[27] Lähnemann D, Köster J, Szczurek E, McCarthy DJ, Hicks SC, Robinson MD, et al. Eleven Grand Challenges in Single-Cell Data Science. Genome Biology. 2020;21(1):31. https://doi.org/10.1186/s13059-020-1926-6.

[28] Laks E, McPherson A, Zahn H, Lai D, Steif A, Brimhall J, et al. Clonal Decomposition and DNA Replication States Defined by Scaled Single-Cell Genome Sequencing. Cell. 2019;179(5):1207–1221.e22. https://doi.org/10.1016/j.cell.2019.10.026.

[29] Navin N, Kendall J, Troge J, Andrews P, Rodgers L, McIndoo J, et al. Tumour Evolution Inferred by Single-Cell Sequencing. Nature. 2011;472(7341):90–94. https://doi.org/10.1038/nature09807.

[30] Wang Y, Waters J, Leung ML, Unruh A, Roh W, Shi X, et al. Clonal Evolution in Breast Cancer Revealed by Single Nucleus Genome Sequencing. Nature. 2014;512(7513):155–160. https://doi.org/10.1038/nature13600.

[31] Zahn H, Steif A, Laks E, Eirew P, VanInsberghe M, Shah SP, et al. Scalable Whole-Genome Single-Cell Library Preparation without Preamplification. Nature Methods. 2017;14(2):167–173. https://doi.org/10.1038/nmeth.4140.

[32] Vitak SA, Torkenczy KA, Rosenkrantz JL, Fields AJ, Christiansen L, Wong MH, et al. Sequencing Thousands of Single-Cell Genomes with Combinatorial Indexing. Nature Methods. 2017;14(3):302–308. https://doi.org/10.1038/nmeth.4154.

[33] Mulqueen RM, Pokholok D, O’Connell BL, Thornton CA, Zhang F, O’Roak BJ, et al. High-Content Single-Cell Combinatorial Indexing. Nature Biotechnology. 2021;39(12):1574–1580. https://doi.org/10.1038/s41587-021-00962-z.

[34] Minussi DC, Nicholson MD, Ye H, Davis A, Wang K, Baker T, et al. Breast Tumours Maintain a Reservoir of Subclonal Diversity during Expansion. Nature. 2021;592(7853):302–308. https://doi.org/10.1038/s41586-021-03357-x.

[35] 10x Genomics.: Single Cell CNV. https://www.10xgenomics.com/products/single-cell-cnv.

[36] Salcedo A, Tarabichi M, Espiritu SMG, Deshwar AG, David M, Wilson NM, et al. A Community Effort to Create Standards for Evaluating Tumor Subclonal Reconstruction. Nature Biotechnology. 2020;38(1):97–107. https://doi.org/10.1038/s41587-019-0364-z.

[37] Nik-Zainal S, Van Loo P, Wedge DC, Alexandrov LB, Greenman CD, Lau KW, et al. The Life History of 21 Breast Cancers. Cell. 2012;149(5):994–1007. https://doi.org/10.1016/j.cell.2012.04.023.

[38] Jamal-Hanjani M, Wilson GA, McGranahan N, Birkbak NJ, Watkins TBK, Veeriah S, et al. Tracking the Evolution of Non–Small-Cell Lung Cancer. New England Journal of Medicine. 2017;376(22):2109–2121. https://doi.org/10.1056/NEJMoa1616288.

[39] Takahashi S, Miura H, Shibata T, Nagao K, Okumura K, Ogata M, et al. Genome-Wide Stability of the DNA Replication Program in Single Mammalian Cells. Nature Genetics. 2019;51(3):529–540. https://doi.org/10.1038/s41588-019-0347-5.

[40] Van der Aa N, Cheng J, Mateiu L, Esteki MZ, Kumar P, Dimitriadou E, et al. Genome-Wide Copy Number Profiling of Single Cells in S-phase Reveals DNAreplication Domains. Nucleic Acids Research. 2013;41(6):e66–e66. https://doi.org/10.1093/nar/gks1352.

[41] Van Loo P, Nordgard SH, Lingjærde OC, Russnes HG, Rye IH, Sun W, et al. Allele-Specific Copy Number Analysis of Tumors. Proceedings of the National Academy of Sciences. 2010;107(39):16910–16915. https://doi.org/10.1073/pnas.1009843107.

[42] Chen H, Bell JM, Zavala NA, Ji HP, Zhang NR. Allele-Specific Copy Number Profiling by next-Generation DNA Sequencing. Nucleic Acids Research. 2015;43(4):e23. https://doi.org/10.1093/nar/gku1252.

[43] Favero F, Joshi T, Marquard AM, Birkbak NJ, Krzystanek M, Li Q, et al. Sequenza: Allele-Specific Copy Number and Mutation Profiles from Tumor Sequencing Data. Annals of Oncology. 2015;26(1):64–70. https://doi.org/10.1093/annonc/mdu479.

[44] Shen R, Seshan VE. FACETS: Allele-Specific Copy Number and Clonal Heterogeneity Analysis Tool for High-Throughput DNA Sequencing. Nucleic Acids Research. 2016;44(16):e131. https://doi.org/10.1093/nar/gkw520.

[45] Cun Y, Yang TP, Achter V, Lang U, Peifer M. Copy-Number Analysis and Inference of Subclonal Populations in Cancer Genomes Using Sclust. Nature Protocols. 2018;13(6):1488–1501. https://doi.org/10.1038/nprot.2018.033.

[46] Poell JB, Mendeville M, Sie D, Brink A, Brakenhoff RH, Ylstra B. ACE: Absolute Copy Number Estimation from Low-Coverage Whole-Genome Sequencing Data. Bioinformatics. 2019;35(16):2847–2849. https://doi.org/10.1093/bioinformatics/bty1055.

[47] Sauer CM, Eldridge MD, Vias M, Hall JA, Boyle S, Macintyre G, et al. Absolute Copy Number Fitting from Shallow Whole Genome Sequencing Data. bioRxiv. 2021;p. 2021.07.19.452658. https://doi.org/10.1101/2021.07.19.452658.

[48] Zaccaria S, Raphael BJ. Accurate Quantification of Copy-Number Aberrations and Whole-Genome Duplications in Multi-Sample Tumor Sequencing Data. Nature Communications. 2020;11(1):4301. https://doi.org/10.1038/s41467-020-17967-y.

[49] Mallory XF, Edrisi M, Navin N, Nakhleh L. Assessing the Performance of Methods for Copy Number Aberration Detection from Single-Cell DNA Sequencing Data. PLOS Computational Biology. 2020;16(7):e1008012. https://doi.org/10.1371/journal.pcbi.1008012.

[50] Lai D, Ha G, Shah S.: Package ‘HMMcopy’.

[51] Knouse KA, Wu J, Amon A. Assessment of Megabase-Scale Somatic Copy Number Variation Using Single-Cell Sequencing. Genome Research. 2016;26(3):376–384. https://doi.org/10.1101/gr.198937.115.

[52] Funnell T, O’Flanagan CH, Williams MJ, McPherson A, McKinney S, Kabeer F, et al. Single-Cell Genomic Variation Induced by Mutational Processes in Cancer. Nature. 2022;612(7938):106–115. https://doi.org/10.1038/s41586-022-05249-0.

[53] Markowska M, Caka-la T, Miasojedow B, Aybey B, Juraeva D, Mazur J, et al. CONET: Copy Number Event Tree Model of Evolutionary Tumor History for Single-Cell Data. Genome Biology. 2022;23(1):128. https://doi.org/10.1186/s13059-022-02693-z.

[54] Salehi S, Dorri F, Chern K, Kabeer F, Rusk N, Funnell T, et al. Cancer Phylogenetic Tree Inference at Scale from 1000s of Single Cell Genomes. bioRxiv. 2023;p. 2020.05.06.058180. https://doi.org/10.1101/2020.05.06.058180.

[55] Garvin T, Aboukhalil R, Kendall J, Baslan T, Atwal GS, Hicks J, et al. Interactive Analysis and Assessment of Single-Cell Copy-Number Variations. Nature Methods. 2015;12(11):1058–1060. https://doi.org/10.1038/nmeth.3578.

[56] Mallory XF, Edrisi M, Navin N, Nakhleh L. Methods for Copy Number Aberration Detection from Single-Cell DNA-sequencing Data. Genome Biology. 2020;21(1):208. https://doi.org/10.1186/s13059-020-02119-8.

[57] Wang R, Lin DY, Jiang Y. SCOPE: A Normalization and Copy-Number Estimation Method for Single-Cell DNA Sequencing. Cell Systems. 2020;10(5):445– 452.e6. https://doi.org/10.1016/j.cels.2020.03.005.

[58] Zaccaria S, Raphael BJ. Characterizing Allele- and Haplotype-Specific Copy Numbers in Single Cells with CHISEL. Nature Biotechnology. 2021;39(2):207–214. https://doi.org/10.1038/s41587-020-0661-6.

[59] Andor N, Lau BT, Catalanotti C, Sathe A, Kubit M, Chen J, et al. Joint Single Cell DNA-seq and RNA-seq of Gastric Cancer Cell Lines Reveals Rules of in Vitro Evolution. NAR Genomics and Bioinformatics. 2020;2(2). https://doi.org/10.1093/nargab/lqaa016.

[60] Weiner AC, Williams MJ, Shi H, Shah SP, McPherson A. Modeling Single Cell DNA Replication Dynamics and Aneuploidy in Genomically Unstable Cancers. bioRxiv. 2023;p. 2023.04.10.536250. https://doi.org/10.1101/2023.04.10.536250.

[61] Killick R, Fearnhead P, Eckley IA. Optimal Detection of Changepoints With a Linear Computational Cost. Journal of the American Statistical Association. 2012;107(500):1590–1598. https://doi.org/10.1080/01621459.2012.737745.arxiv:1101.1438.

[62] Hipel KW, McLeod AI. Time Series Modelling of Water Resources and Environmental Systems. Elsevier; 1994.

[63] Taliun D, Harris DN, Kessler MD, Carlson J, Szpiech ZA, Torres R, et al. Sequencing of 53,831 Diverse Genomes from the NHLBI TOPMed Program. Nature. 2021;590(7845):290–299. https://doi.org/10.1038/s41586-021-03205-y.

[64] Anscombe FJ. Sampling Theory Of The Negative Binomial And Logarithmic Series Distributions. Biometrika. 1950;37(3-4):358–382. https://doi.org/10.1093/biomet/37.3-4.358.

[65] Bliss CI, Fisher RA. Fitting the Negative Binomial Distribution to Biological Data. Biometrics. 1953;9(2):176–200. https://doi.org/10.2307/3001850. 3001850.

[66] Anraku K, Yanagimoto T. Estimation for the Negative Binomial Distribution Based on the Conditional Likelihood. Communications in Statistics - Simulation and Computation. 1990;19(3):771–786. https://doi.org/10.1080/03610919008812887.

[67] Blei DM, Jordan MI. Variational Inference for Dirichlet Process Mixtures. Bayesian Analysis. 2006;1(1):121–143. https://doi.org/10.1214/06-BA104.

[68] Hoffman MD, Blei DM, Wang C, Paisley J. Stochastic Variational Inference. The Journal of Machine Learning Research. 2013;14(1):1303–1347.

[69] Abadi M, Agarwal A, Barham P, Brevdo E, Chen Z, Citro C, et al.: TensorFlow: Large-scale Machine Learning on Heterogeneous Systems.

[70] Hansen RS, Thomas S, Sandstrom R, Canfield TK, Thurman RE, Weaver M, et al. Sequencing Newly Replicated DNA Reveals Widespread Plasticity in Human Replication Timing. Proceedings of the National Academy of Sciences. 2010;107(1):139–144. https://doi.org/10.1073/pnas.0912402107.

[71] Thurman RE, Day N, Noble WS, Stamatoyannopoulos JA. Identification of Higher-Order Functional Domains in the Human ENCODE Regions. Genome Research. 2007;17(6):917–927. https://doi.org/10.1101/gr.6081407.

[72] Scheinin I, Sie D, Bengtsson H, van de Wiel MA, Olshen AB, van Thuijl HF, et al. DNA Copy Number Analysis of Fresh and Formalin-Fixed Specimens by Shallow Whole-Genome Sequencing with Identification and Exclusion of Problematic Regions in the Genome Assembly. Genome Research. 2014;24(12):2022–2032. https://doi.org/10.1101/gr.175141.114.

[73] Langdon SP, Lawrie SS, Hay FG, Hawkes MM, McDonald A, Hayward IP, et al. Characterization and Properties of Nine Human Ovarian Adenocarcinoma Cell Lines. Cancer Research. 1988;48(21):6166–6172.

[74] Cooke SL, Ng CKY, Melnyk N, Garcia MJ, Hardcastle T, Temple J, et al. Genomic Analysis of Genetic Heterogeneity and Evolution in High-Grade Serous Ovarian Carcinoma. Oncogene. 2010;29(35):4905–4913. https://doi.org/10.1038/onc.2010.245.

[75] Sakai W, Swisher EM, Jacquemont C, Chandramohan KV, Couch FJ, Langdon SP, et al. Functional Restoration of BRCA2 Protein by Secondary BRCA2 Mutations in BRCA2-Mutated Ovarian Carcinoma. Cancer Research. 2009;69(16):6381–6386. https://doi.org/10.1158/0008-5472.CAN-09-1178.

[76] Sakaue-Sawano A, Kurokawa H, Morimura T, Hanyu A, Hama H, Osawa H, et al. Visualizing Spatiotemporal Dynamics of Multicellular Cell-Cycle Progression. Cell. 2008;132(3):487–498. https://doi.org/10.1016/j.cell.2007.12.033.

[77] Koh SB, Mascalchi P, Rodriguez E, Lin Y, Jodrell DI, Richards FM, et al. A Quantitative FastFUCCI Assay Defines Cell Cycle Dynamics at a Single-Cell Level. Journal of Cell Science. 2017;130(2):512–520. https://doi.org/10.1242/jcs.195164.

[78] Van Oudenhove EPK. Integrative Assessment of Homologous Recombination Deficiency in Ovarian Cancer. Cambridge: University of Cambridge; 2016.

[79] Darzynkiewicz Z, Juan G, Bedner E. Determining Cell Cycle Stages by Flow Cytometry. Current Protocols in Cell Biology. 1999;1(1):8.4.1–8.4.18. https://doi.org/10.1002/0471143030.cb0804s01.

[80] Loh PR, Danecek P, Palamara PF, Fuchsberger C, A Reshef Y, K Finucane H, et al. Reference-Based Phasing Using the Haplotype Reference Consortium Panel. Nature Genetics. 2016;48(11):1443–1448. https://doi.org/10.1038/ng.3679.

